# YAP localization mediates mechanical adaptation of human cancer cells during extravasation *in vivo*

**DOI:** 10.1101/2023.11.14.567015

**Authors:** Woong Young So, Claudia S. Wong, Udochi F. Azubuike, Colin D. Paul, Paniz Rezvan Sangsari, Patricia B. Gordon, Hyeyeon Gong, Tapan K. Maity, Perry Lim, Zhilin Yang, Christian A. Haryanto, Eric Batchelor, Lisa M. Jenkins, Nicole Y. Morgan, Kandice Tanner

**Author notes:** Corresponding author information: Dr. Kandice Tanner; Center for Cancer Research, National Cancer Institute, Building 37, Room 2132, Bethesda, MD 20892.

## Abstract

Biophysical profiling of primary tumors has revealed that individual tumor cells fall along a highly heterogeneous continuum of mechanical phenotypes. One idea is that a subset of tumor cells is “softer” to facilitate detachment and escape from the primary site, a step required to initiate metastasis. However, it has also been postulated that cells must be able to deform and generate sufficient force to exit into distant sites. Here, we aimed to dissect the mechanical changes that occur during extravasation and organ colonization. Using multiplexed methods of intravital microscopy and optical tweezer based active microrheology, we obtained longitudinal images and mechanical profiles of cells during organ colonization *in vivo*. We determined that cells were softer, more liquid like upon exit of the vasculature but stiffened and became more solid like once in the new organ microenvironment. We also determined that a YAP mediated mechanogenotype influenced the global dissemination in our in vivo and in vitro models and that reducing mechanical heterogeneity could reduce extravasation. Moreover, our high throughput analysis of mechanical phenotypes of patient samples revealed that this mechanics was in part regulated by the external hydrodynamic forces that the cancer cells experienced within capillary mimetics. Our findings indicate that disseminated cancer cells can keep mutating with a continuum landscape of mechano-phenotypes, governed by the YAP-mediated mechanosensing of hydrodynamic flow.

## Introduction

The metastatic process encompasses the escape, survival, adaptation and proliferation of tumor cells at a distal site[1–3]. Seeding of secondary sites is an inefficient process where only 0.01% of the disseminated tumor cells (DTCs) result in malignant outgrowth[2, 4]. Low survival of DTCs is in part due to the environmental cues such as hemodynamic forces, immune surveillance and anoikis that these cells experience during transit[2, 4]. In spite of this metastatic inefficiency, this advanced stage of cancer remains the main root of lethality[1, 3]. One possible curative intervention is employing therapeutics that directly target these errant DTCs such that they are eliminated to avoid metastatic disease[3, 5]. This approach necessitates a thorough understanding of the mechanisms that regulate the intrinsic or de novo traits of DTCs that enable their survival in the presence of multiple environmental insults. One such “cell-trinsic” factor could be the mechanical phenotype, a key part in how tumor cells maintain genomic and cytoplasmic integrity under flow induced deformation[6]. Moreover, this factor also plays a role in regulating immune activity via mechanical coupling between cancer cell and immune cells[7–9]. Mechanical phenotype is an umbrella term that incorporates a collection of physical properties including viscoelasticity, cell shape, cell deformability, and adhesion properties[10, 11]. Employing mechanical phenotype as an identifier of extravasation potential necessitates that we understand the biological mechanism that drives these phenotypes in the context of disease.

Curated databases have identified differential expression, mutational and alternatively spliced gene expression levels of a myriad of genes in cancer cells compared to that measured for normal matched tissues[12–14]. One major thrust in the field of cancer mechanobiology is to link these transcriptomic changes to cancer cell mechanical phenotypes[15–17]. One challenge is that genes identified in comparative arrays can show pleiotropic effects on processes such as proliferation, migration and cell adhesion, which are all important along the continuum that is the metastatic cascade[18, 19]. Moreover, their function in normal homeostasis can be dissimilar from their function in the context of disease and hence will not predict pathologic function[20–22]. Dissecting how these intertwined and dynamic characteristics are modulated along the metastatic cascade are prerequisites in facilitating our understanding of this complex problem.

Interrogation of the complexity of metastasis requires a model system that recreates physiologically relevant environmental dynamics concomitant with the ability to visualize and mechanically profile the tumor cells for the latter stages of metastasis[1, 23]. Here, we propose that multiplexed approaches are needed to bring insight to this important question. Direct examination of each of the discrete steps of the metastatic cascade remains challenging in models of metastasis such as murine models[1, 23]. Moreover, coupling imaging techniques with that of functional modalities such as single cell mechanical profiling *in vivo* adds a level of difficulty that is rarely achieved for the latter stages of the metastatic cascade. Recently, the zebrafish model has been employed to recreate intravasation, vascular transit, extravasation and survival in distant organs[24–28]. This model has also been amenable for probing the role of biophysical properties during the latter stages of metastasis[29, 30]. Moreover, these studies have been instrumental in elucidating the role of biophysical properties of the microenvironment in driving vascular transit and organ specific metastasis[24, 25]. However, resolution of how the dynamic coupling between the tumor cells and environmental cues direct cell fate decisions from transit to exit and survival in a distant organ is needed.

Focusing on disseminated tumor cells, we previously employed the human xenograft in zebrafish model to recreate the fluid dynamics experienced as cells move within, occlude and exit capillary beds as first steps of organ colonization. Of all the organs, the brain remains the most technically challenging for visualizing the earliest stages of metastasis.[31] Due to the ease of longitudinal imaging in the zebrafish model, in this work we go a step further by looking at the nascent stages of organ colonization one day after extravasation into brain parenchyma. We apply multiplexed methods of intravital microscopy and optical tweezer based active microrheology to combine longitudinal images and mechanical profiles of cells *in vivo*. We complement our studies with *in vitro* models that recreated tumor cell movement in confined spaces and methodologies that provided a high throughput analysis of single cell mechanical phenotypes under variable flows. Using this combinatorial approach, we determine that an emergent and dynamic mechanical phenotype facilitated exit, entry and adaptation into the brain microenvironment. We then aimed to understand a mechanogenetic factor that regulates this process. YAP, a transcriptional coactivator, touted as a mechanically sensitive transcription factor, also played a role in regulating the tumor cells’ ability to tune their mechanical phenotype thus altering the extravasation potential. Our data provides evidence for a link between mechanically mediated extravasation potential and activity of a transcription factor that has also been observed to be upregulated in many human cancers.

## Results

### Occluded cells in the capillary bed have distinct viscoelastic properties from that measured for extravasated cells *in vivo*

To model transit of circulating tumor cells through the vasculature, we injected human breast tumor and melanoma cells pre-labeled with micron diameter beads directly into the circulation of larval zebrafish (Figure 1a). After one day post injection, we visualized cancer cells that have traveled and remain immobilized within blood vessels, and those that had successfully extravasated and entered the brain (Figure 1b and Supplemental movie 1). During extravasation, it is thought that cells deform and alter mechanical properties to breach the endothelial barrier[32, 33]. Thus, we designed a microfluidic device in which cells become occluded at a constriction point prior to deforming and invading through a steric barrier (Figure 1c, Supplemental Figure 1a, and Supplementary movie 2). With these two models in hand, we employed optical tweezer based active microrheology to measure physical changes in the cell’s mechanical phenotype directly *in vivo* and in microfluidic devices. Specifically, we use a vis a vis calibration technique to quantitate the intracellular complex shear modulus, in which the complex modulus, ‘|G*|exp(iδ) = G′(ω) +*i*G ″(ω)’ is composed of the storage (G’(ω) − elastic modulus) and loss moduli (G”(ω) – viscous modulus) (Figure 1d-f)[29, 34–36]. The frequency at which the two curves intersect is defined as the crossover frequency. Shifts in this frequency are used as a metric to determine the relative behavior of either more solid-like (more G’ contribution) or liquid-like (more G” contribution)[37]. Extravasated cells in the brain parenchyma were more liquid-like (lower crossover frequency) than the cells occluded in the intravascular lumenal space (Figure 1d-e). We also performed longitudinal measurement of the same cells immediately and 1 day post extravasation to fully determine the dynamics of the mechanical changes (Figure 1d-e and Supplementary Movie 3) as the extravasated cancer cells were stationary. Over the observation time, the tumor cells modulated their liquid like behaviors: cells during post extravasation became more solid like compared to immediately after exit (Figure 1d-e).

**Figure 1:**
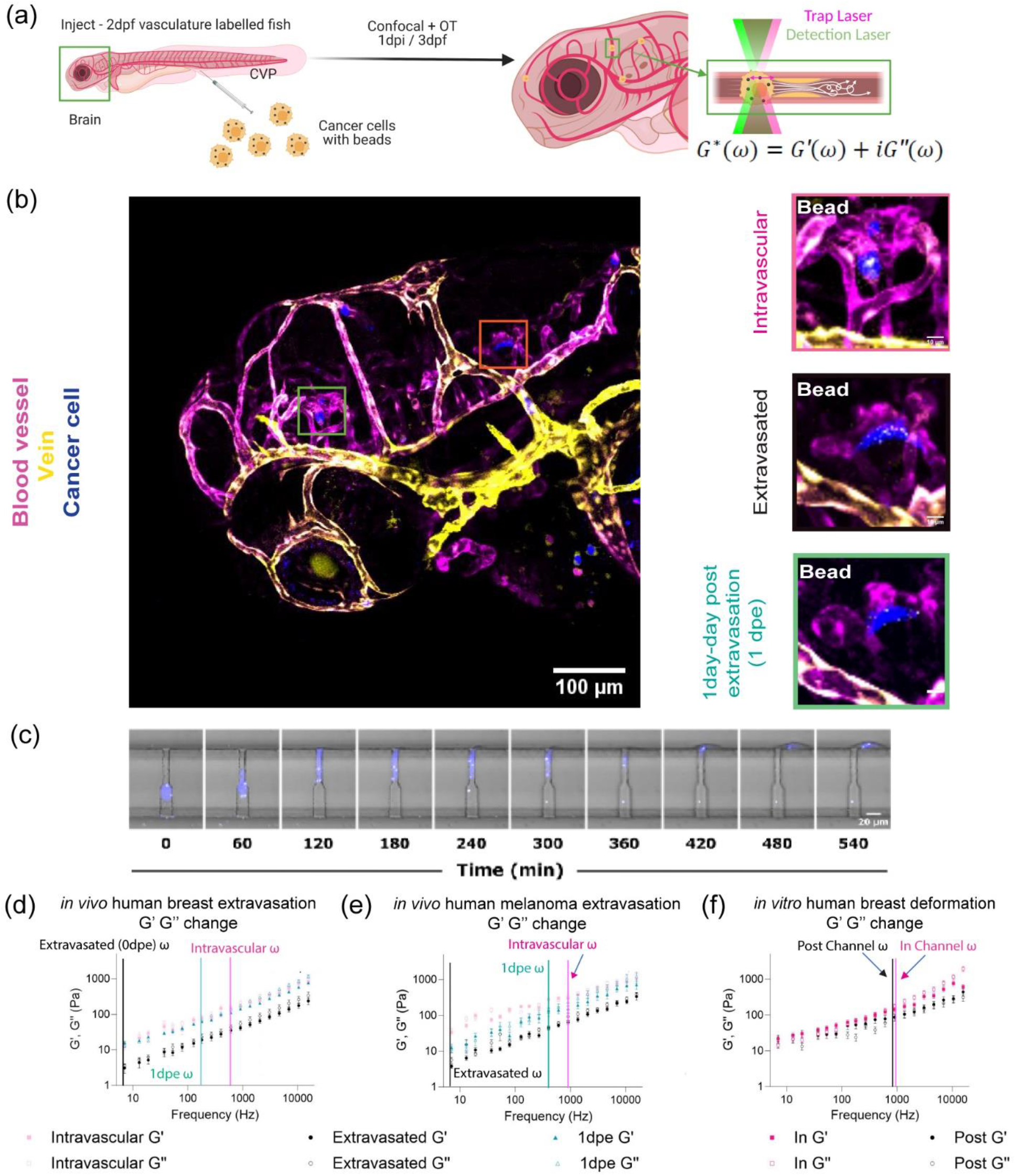
Occluded cells in the capillary bed have distinct viscoelastic properties from that measured for extravasated cells *in vivo.* (a) Scheme of *in vivo* cancer cell mechanical measurement at zebrafish brain by optical trap (OT) where cancer cells with internalized 1 µm beads are injected into the circulation of zebrafish at 2 days-post-fertilization (2dpf) for mechanical mapping at 3dpf and 1 day-post-injection (dpi). (b) Images of capillary arrested (intravascular) and extravasated cancers (blue) inside of zebrafish, Tg(flk:mCherry/MRC1a:EGFP), brain at 3dpf/1dpi along with 1 day-post-extravasated (dpe) from 4dpf/2dpi. (c) Tracking of migration of human breast cancer cell (MDA-MB-231) in extravasation mimicking microfluidics device (confined channel) (d) log-log plot of *in vivo* human breast cancer cell (MDA-MB-231) mechanics (elastic modulus, G’, and viscous modulus, G’’) and frequencies (7Hz to 15kHz) in the function of ‘Intravascular’ (n=22), ‘Extravasated’ (n=5), ‘1dpe’ (n=3). (e) log-log plot of *in vivo* human melanoma (A375SM) mechanics (elastic modulus, G’, and viscous modulus, G’’) and frequencies (7Hz to 15kHz) in the function of ‘Intravascular’ (n=6), ‘Extravasated’ (n=4), ‘1dpe’ (n=3). (f) log-log plot of *in vitro* human breast cancer cell (MDA-MB-231) mechanics (elastic modulus, G’, and viscous modulus, G’’) and frequencies (7Hz to 15kHz) in the function of ‘In Channel’ (n=38) and ‘Post Channel’ (n=10). Error bars in standard of error. Crossover frequency is assigned for each condition to point out the frequency where G’’ becomes more dominant than G’.

As we measured differences in mechanical phenotype, we next asked if this finding was due to mechanical heterogeneity in the population of injected cells, where only cells that were softer can efficiently extravasate, or if dynamic re-organization in response to cues was a key determinant of adaptation required for exit. We next probed cells within the channels vs. those that have migrated through these constrictions smaller than the cell diameter (Figure 1f). Cells within channels were stiffer before successful migration through the steric hindrance and softer after exiting the microchannel (Figure 1f). There was a modest shift in the crossover frequency indicating that the cells were more liquid-like compared to those within the channels (Figure 1f).

### Extravasated cancer cells show comparable mechanical properties after colonization to adapt to the organ microenvironment

The mechanical properties of cells are determined by intracellular networks and the cues received from the microenvironment[21, 23, 38]. We first measured the mechanical properties of the uninjected brain parenchyma (Figure 2a and Supplemental Figure 2a). Next, we compared the intracellular cellular mechanical properties of tumor cells to that of the uninjected brain at 4 days post fertilization (dpf) (Figure 2b-d and Supplemental Figure 1b-c). Comparison of the magnitude of the complex moduli indicated that both the breast and melanoma cells were stiffer one day post extravasation than immediately after extravasation (Figure 2b-c and Supplemental Figure 2b-c). In the case of the breast cancer cells, the magnitudes of the complex moduli recovered to values measured for occluded cells (Figure 2d). The melanoma cells remained softer than the cells that were occluded in the lumenal spaces (Figure 2d). However, values of the magnitude of the complex moduli were comparable in terms of the |G(ω)| one day post extravasation for both types of cancer cells (Figure 2d).We determined that the magnitudes of the complex modulus of the cancer cells one day post extravasation were comparable to the value measured for the uninjected 3dpf brain but softer than the value measured for 4dpf (Figure 2d and Supplemental Figure 2a). Moreover, as the complex moduli followed a power law dependence at high frequencies, we then used this metric to further define the material properties of the cells in terms of polymer physics as a proxy for intracellular cytoskeleton architecture (Figure 2b-c). Power law analysis revealed a shift where occluded cells show an average value of ∼0.51 and ∼0.47 for breast and melanoma cells respectively (more flexible networks) whereas the extravasated cells show an average value of ∼0.63 and ∼0.56 (more semi-flexible networks) but then become more like flexible networks one day post extravasation (Figure 2b-c). We performed similar analysis for the uninjected brain and observed that the brain tissue adopted ∼0.60 and ∼0.57 for brain 3dpf and 4dpf respectively.

**Figure 2:**
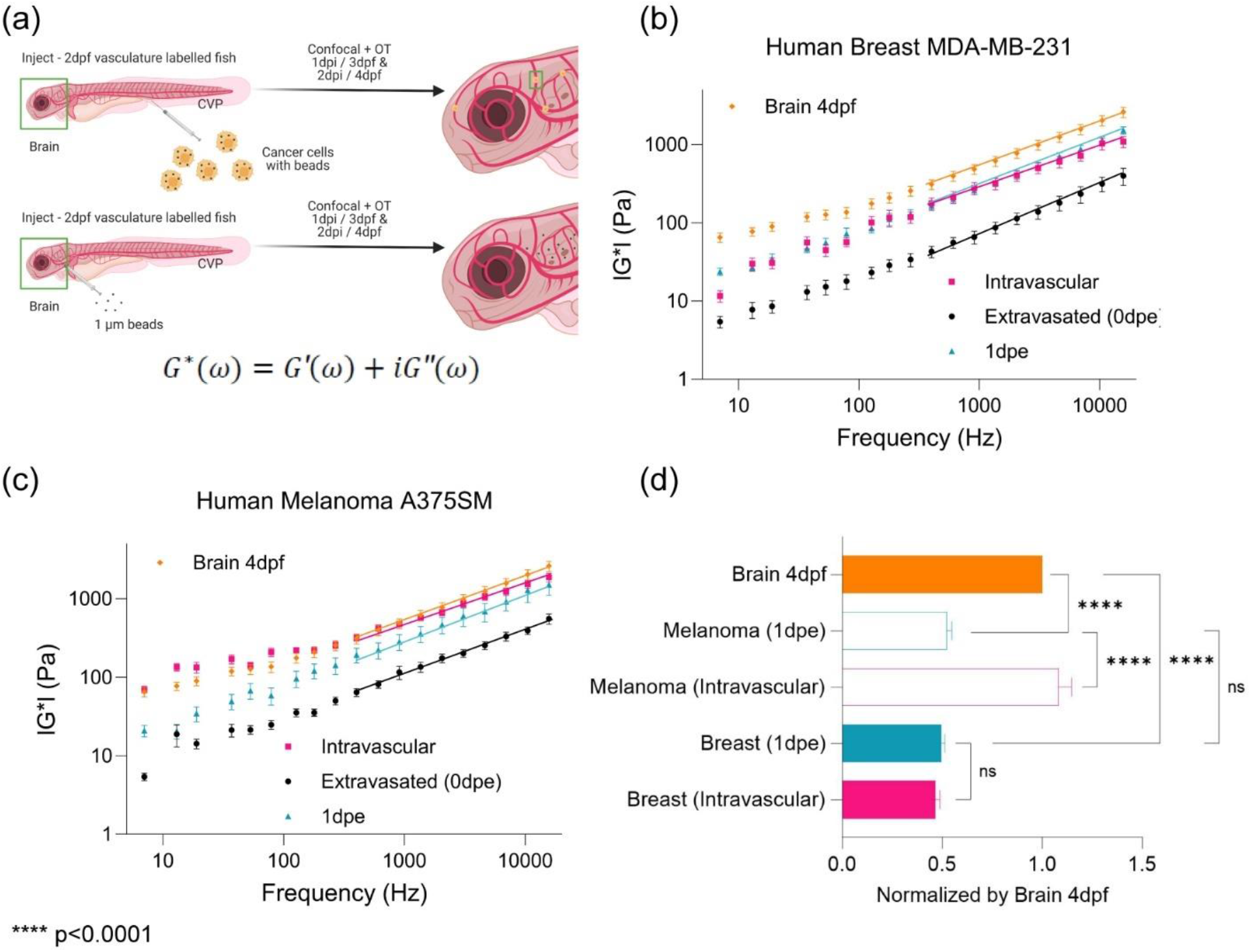
Extravasated cancer cells alter mechanical properties after colonization to adapt to the organ microenvironment. (a) Scheme of following up extravasated cancer cell from 3dpf or 0 day-post-extravasation (dpe) to 4dpf/1dpe inside of zebrafish and of brain mechanics where beads were directly injected into brain of Tg(flk:mCherry/MRC1a:EGFP) at 2dpf for measurement at 3dpf/1dpi and 4dpf/2dpi (b) Complex modulus (|G*|) of human breast cancer cells (MDA-MB-231) in the function of ‘Intravascular’ (n=22), ‘Extravasated (0dpe)’ (n=5), and ‘1dpe’ (n=3) and of brain parenchyma at 4dpf (Number of fish = 5, n=154) (c) Complex modulus (|G*|) of human melanoma cells (A375SM) in the function of ‘Intravascular’ (n=6), ‘Extravasated (0dpe)’ (n=4), and ‘1dpe’ (n=3) and of brain parenchyma at 4dpf (Number of fish = 5, n=154) (d) Normalized bar graph of complex modulus for ‘Breast (Intravascular)’, ‘Breast (1dpe)’, ‘Melanoma (Intravascular)’, ‘Melanoma (1dpe)’, and ‘Brain 4dpf’ in respect to ‘Brain 4dpf’ based on 19 different frequencies from 7Hz to 15kHz. **** p<0.0001, paired two-tailed t-tests.

### Elevated hydrodynamic stresses in capillary size biomimetics drives increased heterogeneity in mechanical phenotypes

During transit, cancer cells experience shear forces due to external blood and lymphatic flows[39, 40]. OT is a low throughput mechanical modality. Thus, to fully investigate how flow may influence the mechanical phenotypes, we employed real-time deformability cytometry (RTDC) to perform high throughput measurements[41–44]. RTDC allows us to mechanically characterize thousands of cells where we can measure both the Young’s modulus (E) – stiffness and deformation (Figure 3a). To simulate different hydrodynamic forces, we modulated the external flow rates to assess the mechanical phenotypes (Supplemental Table 1). We determined that there was a positive correlation between increasing values of average deformation and Young’s modulus as a function of increased flow rates (Figure 3b-c). Examination of the population dynamics revealed that Young’s modulus of the cells adopted a gaussian distribution (Figure 3d-e). We then used the widths of the distribution as a metric of the heterogeneity of the population. The maximum width used as a proxy for maximum heterogeneity was observed at the highest flow rate whereas reduced heterogeneity was observed for the slowest flow rate (Figure 3d-e). We next observed the relationship between deformation and Young’s modulus where deformation and Young’s modulus have an inverse relationship (Supplemental Figure 3a-b).

**Figure 3:**
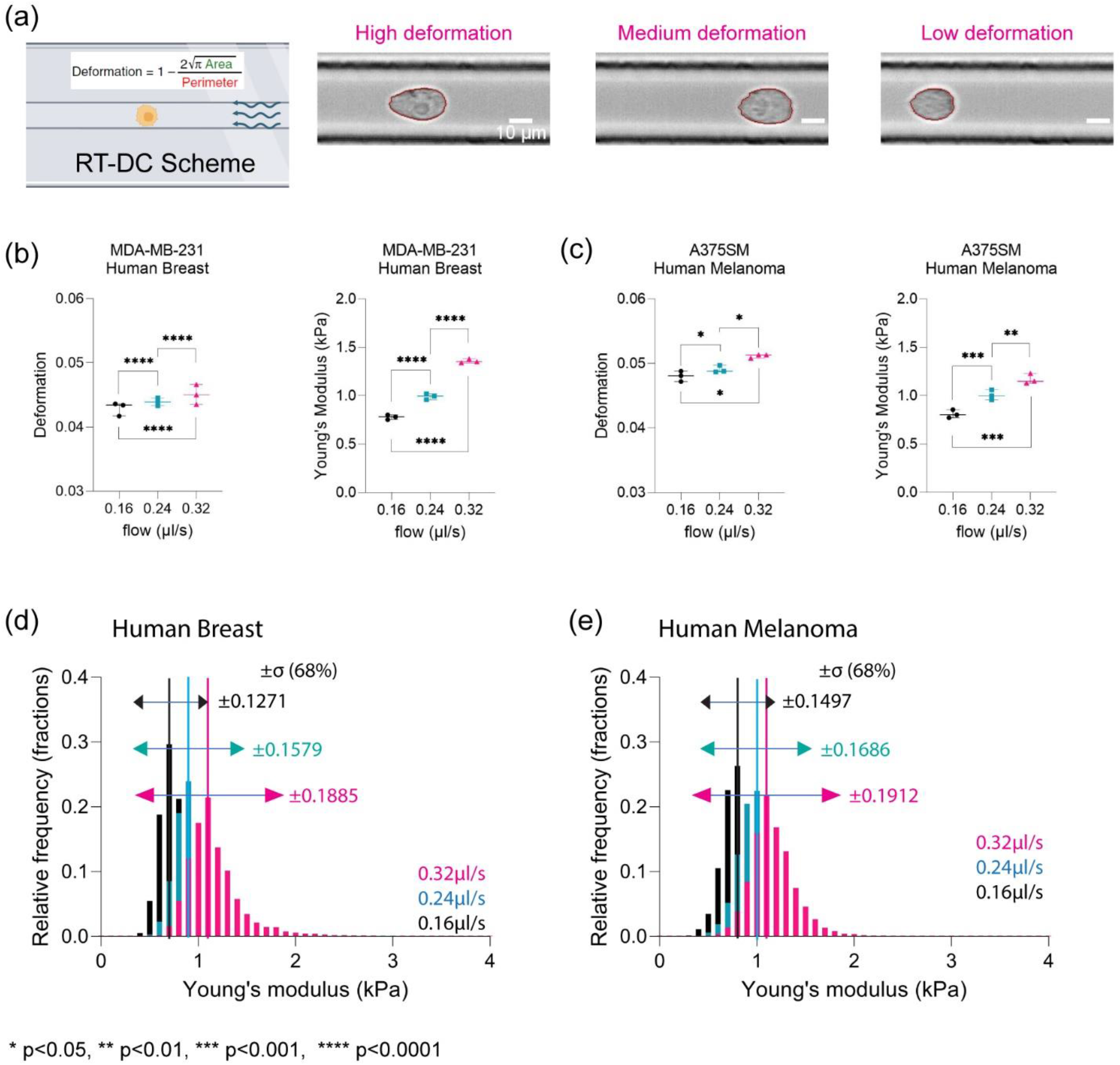
An increase in hydrodynamic stresses in capillary size biomimetics drives increased heterogeneity in emergent size dependent mechanical phenotypes. (a) Scheme of Real-Time Deformability Cytometry (RTDC) measuring physical characteristics of cells under various flow velocities. Examples of cells with various deformation shapes ranging from low to high deformation. (b) Physical characteristics of human breast cancer cell (MDA-MB-231) in terms of deformation and Young’s modulus (kPa) at 0.16 µl/s (N=3; per each replicate more than 1000 cells), 0.24 µl/s (N=3; per each replicate more than 1000 cells), and 0.32 µl/s (N=3; per each replicate more than 1000 cells). **** p<0.0001, paired two-tailed t-tests. (c) Physical characteristics of human melanoma (A375SM) in terms of deformation and Young’s modulus (kPa) at 0.16 µl/s (N=3; per each replicate more than 1000 cells), 0.24 µl/s (N=3; per each replicate more than 1000 cells), and 0.32 µl/s (N=3; per each replicate more than 1000 cells). * p<0.05, ** p<0.01, *** p<0.001, paired two-tailed t-tests. (d) Normalized histogram of human breast cancer cell for Young’s modulus (kPa) at different flow velocities, 0.16 µl/s (n=4906), 0.24 µl/s (n=7398), and 0.32 µl/s (n=9506), with standard deviation (σ, ±68%) (e) Normalized histogram of human melanoma for Young’s modulus (kPa) at different flow velocities, 0.16 µl/s (n= 3954), 0.24 µl/s (n= 6284), and 0.32 µl/s (n= 8442), with standard deviation (σ, ±68%)

### YAP localization modulated deformation and Youngs moduli for patient samples and immortalized cell lines under elevated hydrodynamic stresses in a capillary mimetic

YAP/TAZ has been implicated in mechanotransduction[45]. Notably, nuclear localization and the associated signaling has been shown to be regulated by applied shear stress[46]. Thus, we explored the role of YAP in mechanical phenotype in the presence of fluid shear forces in the absence and presence of an inhibitor of YAP (Figure 4a). We took advantage of the fact that Verteporfin (YAP inhibitor) is fluorescent when illuminated with 410nm and emission of 690nm to assess persistence of the inhibitory effects[47, 48]. Live cell imaging of cells expressing mCerulean – H2B and a cytoplasmic dye revealed both the dynamics and localization of drug accumulation (Supplemental Figure 4a-c). From these data, we determined that 4 hours treatment was sufficient to maintain a stable phenotype for mechanical mapping. RT DC measurements determined that drug treated cells showed an increased average deformation and reduction in the average stiffnesses as a function of flow rates (Figure 4b, Supplementary Figure 4d-e). Immunofluorescence confirmed increased cytoplasmic localization concomitant with reduced nuclear localization upon treatment with verteporfin (VP) (Figure 4c).

**Figure 4:**
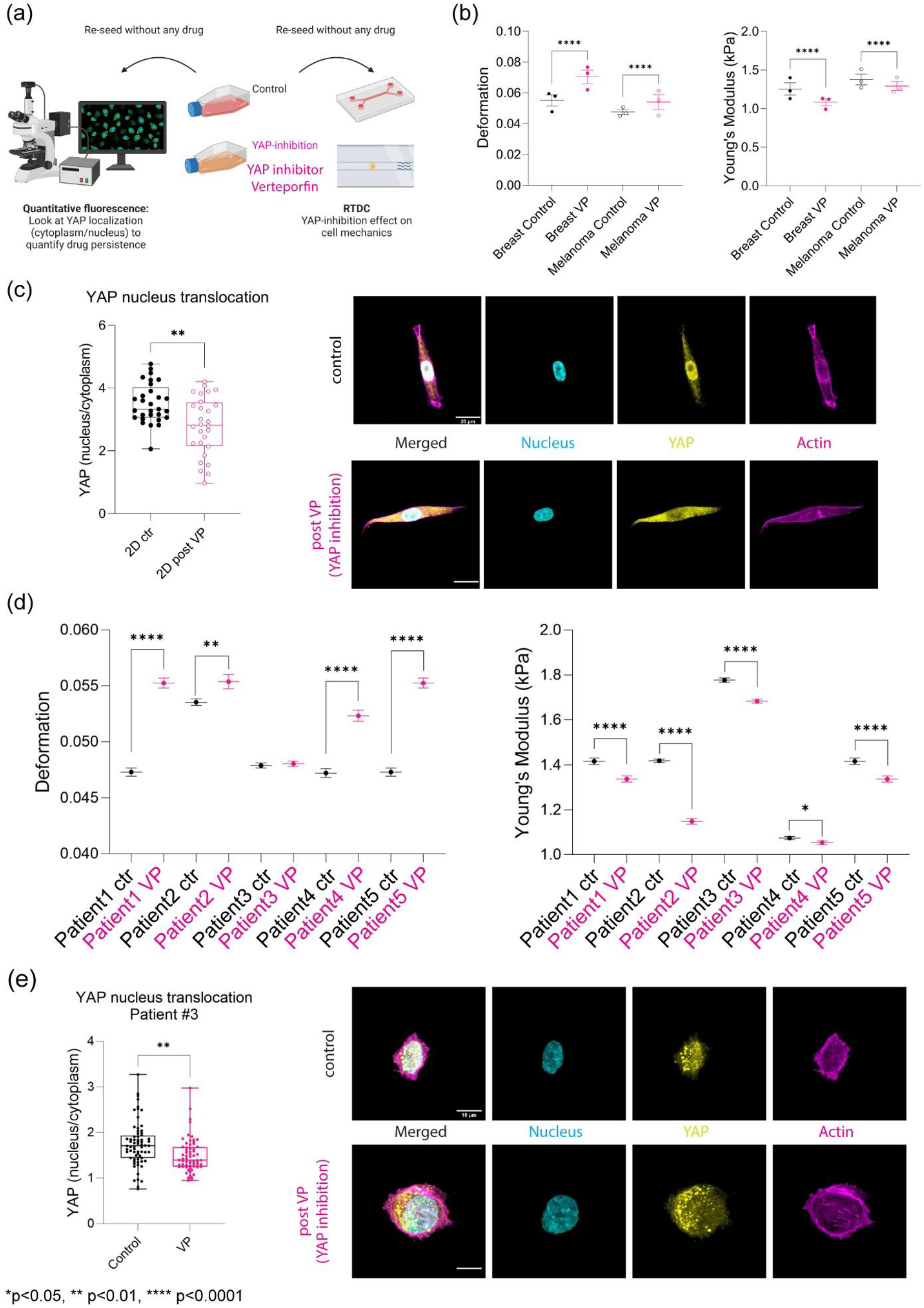
YAP activity modulated deformation and stiffness under fluid stress. (a) Scheme of YAP inhibition using verteporfin (VP), to validate the drug efficacy on human breast cancer cells with quantitative immunofluorescence and RT-DC. (b) Deformation comparison between control/DMSO (N=3; per each replicate more than 1000 cells) and VP (N=3; per each replicate more than 1000 cells) for human breast cancer and human melanoma at 0.32 µl/s. **** p<0.0001, paired two-tailed t-tests. Young’s modulus (kPa) comparison between control/DMSO (N=3; per each replicate more than 1000 cells) and VP (N=3; per each replicate more than 1000 cells) at 0.32 µl/s for human breast cancer and human melanoma at 0.32 µl/s. **** p<0.0001, paired two-tailed t-tests. (c) Quantitative immunofluorescence for human breast cancer cell control/DMSO and post drug treatment (1µg/ml overnight) with nucleus, YAP, and actin. YAP intensity (nucleus/cytoplasm) comparison between control (n=29) and post VP (n=28). ** p<0.01, unpaired two-tailed t-tests. (d) Deformation comparison between control/DMSO and VP at 0.32 µl/s for Patient1 (control n=2667, VP n=2998), Patient2 (control n=5481, VP n=1544), Patient3 (control n=7501, VP n=6330), Patient4 (control n=2132, VP n=1899), and Patient5 (control n=2667, VP n=2998). Error bars are in standard of error. ** p<0.01 and **** p<0.0001, paired two-tailed t-tests. Young’s modulus (kPa) comparison between control/DMSO and VP at 0.32 µl/s for Patient1 (control n=2667, VP n=2998), Patient2 (control n=5481, VP n=1544), Patient3 (control n=7501, VP n=6330), Patient4 (control n=2132, VP n=1899), and Patient5 (control n=2667, VP n=2998). Error bars are in standard of error. * p<0.05 and **** p<0.0001, paired two-tailed t-tests. (e) Quantitative immunofluorescence for Patient3 control/DMSO and post drug treatment (1µg/ml 4-hour treatment) with nucleus, YAP, and actin. YAP intensity (nucleus/cytoplasm) comparison between control (n=78) and post VP (n=79). ** p<0.01, unpaired two-tailed t-tests.

We next probed the role of YAP inhibition on patient samples. Using tumor cells dissociated from patients, we then compared the effect of YAP inhibition on cell deformation and Young’s moduli. We determined that four out of five samples showed a significant increase in deformation upon YAP inhibition whereas all five samples showed reduced Young’s moduli compared to control (Figure 4d). We confirmed that there was an increase in cytoplasmic YAP localization by immunofluorescence of the patient tumor samples upon drug treatment (Figure 4e).

### YAP activity modulated pathways associated with cytoskeletal and nuclear morphology to drive heterogeneity in fluid dependent mechanical phenotypes

Immunoblots demonstrated confirmed reduced activity of YAP and Lamin A and C (Figure 5a-b and Supplementary Figure 4f). Proteomic analysis of whole cell lysates revealed that pathways associated with cell adhesion, cytoskeleton and oxidative phosphorylation were upregulated in control cells compared to the treated cohorts (Figure 5c and Supplementary Figure 4g). Paxillin signaling has been implicated for cancer cell migration and metastasis[49, 50]. On the other hand, pathways associated with integrin signaling and glycolysis metabolism were upregulated in the treated cells compared to the control (Figure 5c and Supplementary Figure 4f). Population dynamics revealed that while the heterogeneity increased as a function of flow rates for both control and VP treated cells (Figure 5d-e and Supplementary Figure 5a-d), the distribution of the YAP inhibited cells was narrower at all flow rates, indicating reduced heterogeneity within the cell population (Figure 5d-e).

**Figure 5:**
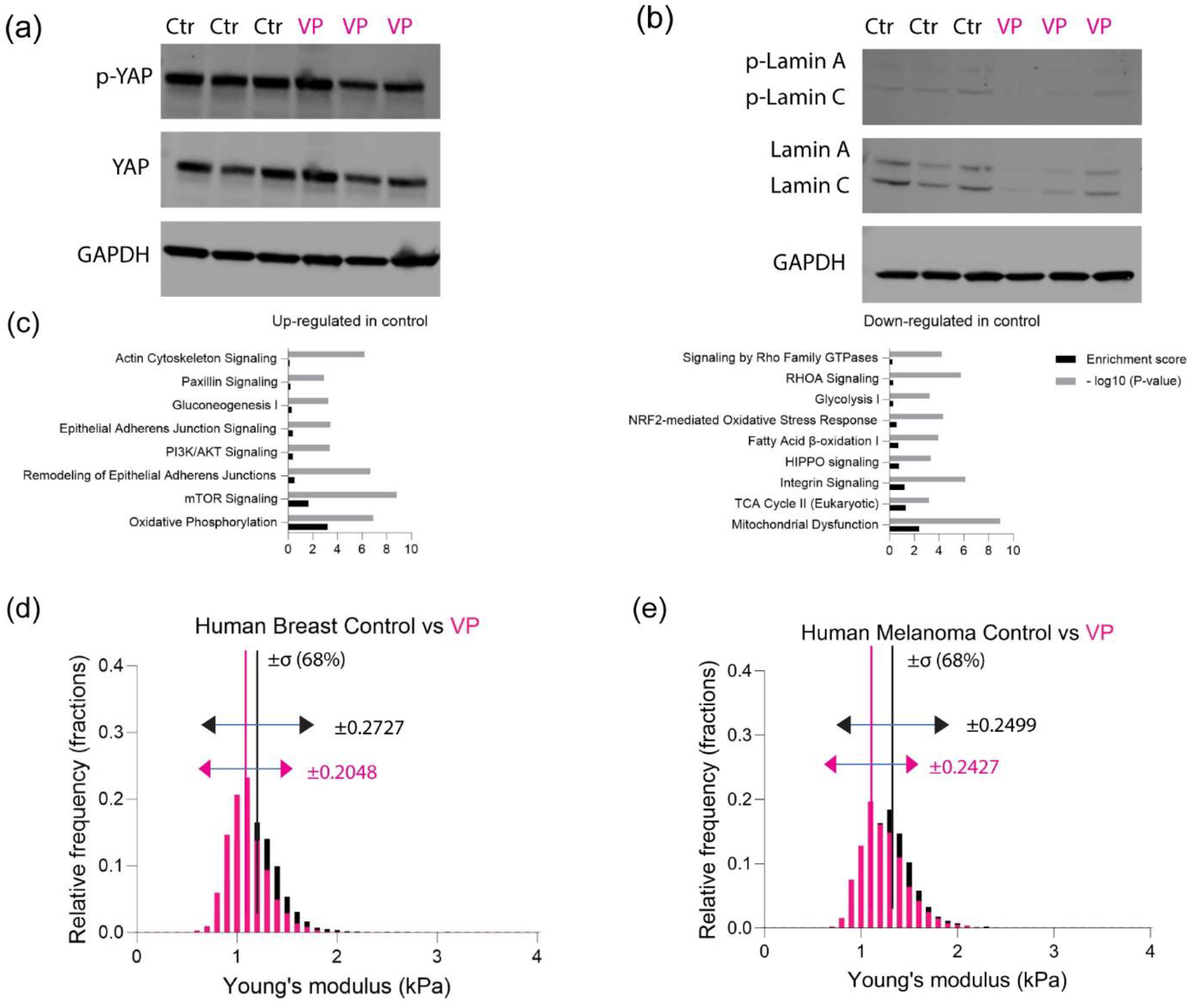
YAP inhibition altered various metabolism and cytoskeleton modeling. (a) Western blot of total YAP with phosphorylated YAP for human breast (N=3) (b) Western blot of total Lamin A/C with phosphorylated Lamin A/C for human breast (N=3) (c) Canonical pathway analysis based on three different replicates of protein lysates analyzed by mass spectrometry, showing up-regulated proteins and down-regulated proteins for human breast control/DMSO. (d) Normalized histogram of human breast cancer cell for young’s modulus (kPa) between control/DMSO and VP at 0.32 µl/s (control n=8712, VP n=10637), with standard deviation (σ, ±68%). (e) Normalized histogram of human melanoma for young’s modulus (kPa) between control/DMSO and VP at 0.32 µl/s (control n=9982, VP n=9988), with standard deviation (σ, ±68%).

**Figure 6:**
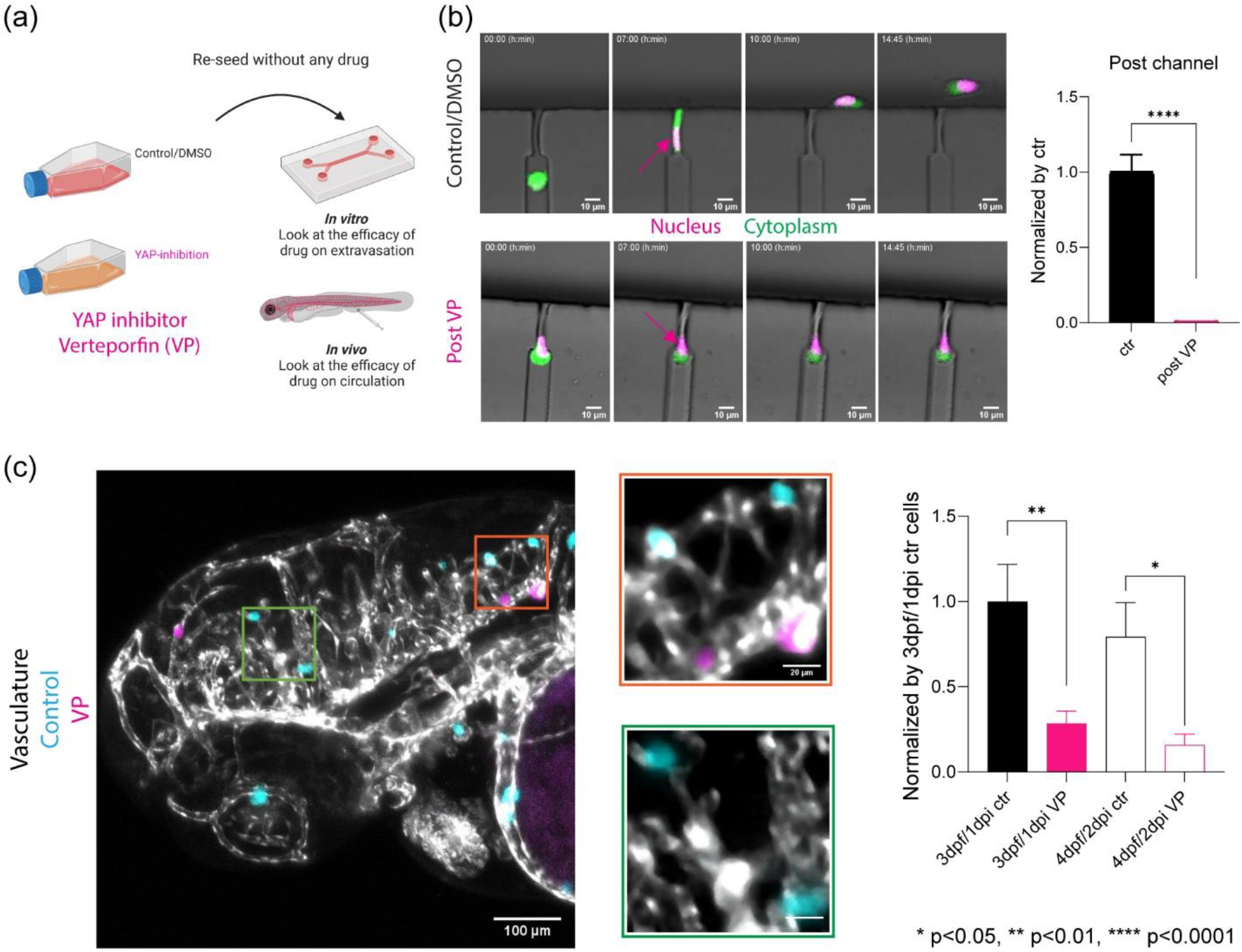
YAP inhibition reduced extravasation in vivo and reduced migration within constrictions in an *in vitro* extravasation device. (a) Scheme for validation of the efficacy of YAP inhibition with verteporfin (VP) on human breast cancer cell extravasation with *in vitro* extravasation mimicking device and *in vivo* zebrafish human cancer cell xenograft. (b) Live cell imaging of control/DMSO cancer cells and VP treated cancer cells with nucleus (H2B) in magenta and membrane dye (CellTracker) in green inside of extravasation mimicking microfluidics devices at different time intervals: Arrows pointing the position of nucleus. Quantification of extravasated cancer cells per device between control/DMSO (n=8) and VP (n=8) after normalization based on control/DMSO. **** p<0.0001, unpaired two-tailed t-tests (c) Zebrafish brain at 3dpf/1dpi after co-injecting with control/DMSO (cyan) and VP (magenta) treated human breast cancer cells into circulation at 2dpf using Tg(flk:mCherry/MRC1a:EGFP); flk and MRC1a are labelled as grey. Quantification of cancer cells at zebrafish brain over time between control/DMSO (3dpf n=18, 4dpf n=8) and VP (3dpf n=18, 4dpf n=8) after normalization based on 3pdf/1dpi control/DMSO. * p<0.05, ** p<0.01, paired two-tailed t-tests.

### YAP inhibition reduced extravasation in vivo and reduced migration in an *in vitro* extravasation device

We next asked if tuning mechanical phenotypes via YAP localization influenced in vivo and in vitro models of extravasation (Figure 5a). As YAP modulation affected the mechanical phenotypes of the cells, we asked if cell behaviors such as confined migration and deformation persist after acute treatment. Verteporfin pre-treated cells were able to migrate within but not escape the constricted spaces (Figure 5b). Direct visualization revealed that the nuclei remained at the front of the cells over the course of ∼14hrs at the impasse compared to vehicle control (Figure 5b and Supplemental Movie 4-5). Following co-injection of control and pre-treated cells into circulation, equal numbers of cells per condition migrated to the brain; however, each population showed distinct differences in extravasation *in vivo* one day and two days post injection. Extravasation of the numbers of pre-treated cells was decreased by 1/3 compared to vehicle control (Figure 5c and Supplemental Movie 6).

## Discussion

The idea of targeting mechanical phenotype as a therapeutic strategy to mitigate metastatic spread has greatly been advanced by the growing appreciation that proteins can be mechanoresponsive. One such protein, YAP/TAZ, is implicated in organ size in normal homeostasis and has also been shown to regulate stem cell differentiation *via* mechanotransduction[45, 51, 52]. YAP has also been shown to promote metastasis in several tumor types by regulating tumor growth[53]. Increased activity of YAP (or its paralog TAZ) has been seen in carcinomas such as breast, lung, liver and pancreatic cancers and neural crest derived cancers such as melanomas and brain cancers[53, 54]. In addition, it is a mechanically sensitive transcription factor, responding to environmental cues such as the physical properties of the extracellular matrix niche and rendering cancer cells insensitive to chemotherapies and targeted therapies[53, 55]. Here, our data adds that YAP plays a pivotal role in how tumor cells respond to fluid forces and navigate within organ capillary beds by regulating mechanical phenotype. Our combined longitudinal intravital imaging with mechanical profiling revealed mechanical adaptation of tumor cells during the initial stages of extravasation and early stages of organ colonization. Moreover, our *in vitro* measurements also quantitated the differential mechanical phenotype during migration in constricted spaces using both high throughput RTDC in capillary mimics and longitudinal cellular imaging and measurement in microfluidics. These observations reinforced that extravasation of tumor cells is an active process and that dynamic mechanical adaptation is vital for progression through the metastatic cascade. It also hints that reducing mechanical heterogeneity could reduce extravasation. These and future studies will be needed to explore if this can be a potential biophysical target in cancer treatment.

Optical tweezers coupled with fluorescence imaging have been employed to quantify mechanical properties of the zebrafish microenvironment for metastasis models highlighting the role of flow in tuning tumor cell arrest[25]. However, understanding the role of physical traits in the exit into distant sites have been restricted to in vitro models of extravasation. Optical tweezers rely on cytoplasmic inclusion of polystyrene beads as sensors to measure microrheological properties of individual cells[37]. These measurements can be used to infer the cells’ mechanical phenotype, which studies have shown is directly related to the coupling of the cytoplasm and nucleus[56]. Other optical methods such as Brillouin microscopy can resolve a proxy of the stiffness of the nucleus and cytoplasm non-invasively[57, 58]. However, our data suggest that additional rheological properties such as hystersivity (solid-like vs liquid-like) are modulated and may be important in understanding organ colonization *in vivo*. This factor concomitant with the high frequency power law dynamics indicates that filamentous components are modulated as a function of organ adaptation[37]. Our proteomic data reinforce that the actin cytoskeleton and focal adhesion signaling pathways are important in regulating the mechanical phenotype. This in concert with recent data expands our repertoire of building a cytoplasmic mechanical fingerprint.

An attractive therapeutic strategy in the arsenal of drug therapeutics addresses survival of disseminated cancer cells as they move within the hematogenous and lymphatic circuitry[59]. The mechanisms of transit can be categorized as active and/ or passive transport[39]. In active transport, transit is influenced by flow but also includes modulation of proteins that regulate adhesion, migration and polarity. These processes are also dynamically regulated by the hydrodynamic forces experienced by the cells. In passive transport, transit is driven solely by hemodynamic or lymphatic forces[39]. However, for each proposed mechanism, the structural integrity of the cells is determined by the robustness of the mechanical phenotype in response to the external stresses[60]. Here, in the experimental regime, flow rates did not induce apoptosis due to rupture such as what is observed for continuous shear stresses above 15 dyn/cm^2^ (1.5 Pa) as the cells pass through the capillary in less than a second. Instead, our high throughput analysis revealed that cells adopted size dependent mechanical phenotypes encoding for deformation and stiffness in response to capillary-hemodynamic forces during transit. Increased heterogeneity in stiffness was observed at higher flow rates and softer cells showed a broader range of deformation. This can have consequences for metastatic spread as cells will experience differential flow rates as they traverse different arms of the circulation system, implying that the route of transit will drive differential mechanical phenotypes that can in turn influence exit into distant sites.

Cells experience an average shear stress of 0.64 dyn/cm^2^ (0.064 Pa) continuously in the lymphatic system which is orders of magnitude less than in the hematogenous network, where the average shear stress ranges from 4–70 dyn/cm^2^ (0.4 – 7 Pa) in the arterial and 1–6 dyn/cm^2^ (0.1 – 0.6 Pa) in the venous systems respectively[61]. It may explain why some cancers preferentially employ lymphatic vs. hematogenous exits. Moreover, the local shear fluid forces may select for hot spots of extravasation due to the multitude of emergent mechanical states. Thus, in the case where cells adopted a more uniform mechanical phenotype in response to lower flow may bias exit at a given site. However, there are important differences to note that blood flow in the zebrafish model is around 0.3 – 2.5 mm/s which is comparable to human and other mammalian systems[40]. Shear forces will vary on many factors including the diameter of the blood vessel[39]. Here, the size of the blood vessels are comparable to sizes observed in capillary beds[24]. Here with RTDC measurements, the cancer cells acutely experience up to ∼18000 dyn/cm^2^ (∼1800 Pa) during transit in the device as the cells are in the capillary less than a second thereby avoiding apoptosis [41–44].

We observed that inhibition of YAP also regulated key aspects of the Lamin machinery where there was a reduction in Lamin a/c activity. Lamin a/c has been implicated in mechanical integrity of the nucleus[56, 62–64]. In one study, mouse embryonic fibroblasts from Lamin a/c –/-mice show a reduction in nuclear stiffness concomitant with increased nuclear fragility resulting in increased apoptosis when these cells experienced mechanical strain[62]. As previously mentioned, we did not observe an increase in nuclear rupture possibly due to a reduction of Lamin a/c activity. Simply, the residual activity and not elimination might be sufficient to affect rigidity but might guard against nuclear rupture.

In a zebrafish model of human cancer cells extravasation, enhanced metastatic spread was observed for tumor cells with constitutively active YAP[65]. Instead of becoming occluded in the first capillary bed, the modified cancer cells were able to re-enter circulation, resulting in increased extravasation events in the brain[65]. Our observations that YAP mediates emergent mechanical phenotypes provide one explanation. Here, we observed that a population of thousands of cells with reduced YAP activity were more uniformly softer and showed increased deformation compared to a comparable number of control cells. The observation that cancer cells were able to migrate but not exit through a constriction or endothelial barrier suggest that while the cells are more deformable in concert with downregulation of cytoskeletal machinery a minimum stiffness may be needed to propel cells across steric hindrances. Moreover, the different localization of the nucleus front vs rear during exit for YAP inhibition vs control may be similar to what has been observed for pressure mediated “piston migration”, which is regulated in part by a nucleoskeleton-intermediate filament linker protein and actomyosin contractility[66, 67]. It cautions us in that spatial localization of organelles such as the nucleus might be needed to interpret single cell mechanical mapping in regulating migration across different tissues. Finally, our proteomic data also highlighted differences in pathways associated with glycolysis and oxidative phosphorylation which are key aspects of tumor metabolism. YAP proteins promote metabolism by regulating glucose uptake and insulin signaling[68, 69]. It remains to be seen if there is indeed a link between metabolic pathways and the mechanical fingerprint.

## Acknowledgements

This effort was supported by the Intramural Research Program of the National Institutes of Health, the National Cancer Institute.

## Methods

### Zebrafish Husbandry

All animal experiments were done under protocols approved by the National Cancer Institute (NCI) and the National Institutes of Health (NIH) Animal Care and Use Committee. Zebrafish were maintained at 28.5 °C on a 14-hour light/10-hour dark cycle according to standard procedures. Embryos were obtained from natural spawning, raised at 28.5 °C, and maintained in fish water, 60 mg sea salt (Instant Ocean, #SS15-10) per liter of DI water. For all experiments, larvae were transferred to fish water supplemented with N-phenylthiourea (PTU; Millipore Sigma, # P7629-25G) between 18-22 hours post-fertilization to inhibit melanin formation for enhanced optical transparency. PTU fish embryo water was prepared by dissolving 400 µl of PTU stock (7.5% w/v in DMSO) per 1 L of fish water. Water was replaced twice per day. Embryos were then returned to the incubator at 28.5 °C and checked for normal development. Tg(flk:mCherry/MRC1a:EGFP) zebrafish in Casper background were used to study the mechanics of cancer cells during extravasation at 3 and 4 dpf; eGFP is expressed under the MRC1a promoter to label veins while mCherry is expressed under flk promoter to visualize blood vessels. At 2 dpf, zebrafish were dechorionated using 2 mg/ml pronase (Millipore Sigma, #10165921001) in PTU fish water for injection into the caudal vein of the zebrafish to deliver cells directly into the circulation.

### Cell Culture and Preparation for xenografts in zebrafish

MDA-MB-231 human breast adenocarcinoma cells and A375SM human melanoma cells were grown and maintained in high glucose (4.5 g/l) DMEM containing 10% FBS, 1% L-glut, and 1% P/S at 37 °C and 5% CO_2_ as previously described[24, 70, 71]. To establish MDA-MB-231 and A375SM cell lines expressing H2B-mCerulean3, cells were transduced with lentivirus expressing pLentiPGK Hygro DEST H2B-mCerulean3 (Addgene: 90234) diluted 1:50 in growth medium supplemented with 8 μg/ml protamine sulfate and 10 μM HEPES. After 72 hours, media was removed and selection was performed in growth medium supplemented with 500 μg/ml hygromycin[72]. Positive cells were isolated *via* Fluorescence-Activated Cell Sorting (FACS). Expression of H2B-mCerulean was confirmed by fluorescence microscopy. For circulation injection preparation, cells were detached with 10 mM EDTA in PBS for 5 min at 37 °C, resuspended in growth medium, spun down at 1000 rpm for 5 minutes. Pelleted cells were resuspended in growth medium again for counting. Cells were again centrifuged at 1000 rpm for 5 minutes and resuspended at a concentration of 1×10^6^ cells/ml in PBS containing CellTracker Deep Red (Thermo Fisher Scientific, #C34565) at 1 mM or CellTracker Blue CMAC (Thermo Fisher Scientific, #C2210) at 10 mM. Cells were then labeled with CellTracker at 37 °C for 30 minutes. Labeled cells were washed with PBS twice, with spins at 1000 rpm for 3 minutes between washes. After the final spin, cells were resuspended at 1.5 × 10^6^ cells/mL in growth medium to mix with blue fluorescence 1 μm diameter polystyrene beads (Thermo Fisher Scientific, #F13080) at a ratio of 50 μL stock beads per 500 μL cell suspension. Cells were incubated with beads for 20 minutes at 37 °C. After the incubation, cell-bead mixture was washed twice with PBS, with spins at 1000 rpm for 5 minutes between washes to remove excess beads. After the final spin, cells were resuspended to a concentration of 1 million cells/20 µl in PBS. Cells were kept on ice for 2-3 hours during injection.

Tricaine stock was prepared by dissolving 400 mg of Tricaine powder (ethyl 3-aminobenzoate methanesulfonate; Millipore Sigma, #E10521-50G) with 97.9 ml of deionized water and 2.1 ml of 1 M Tris. Anesthetic fish water was prepared by mixing 4.2 ml of tricaine stock per 100 ml of fish water supplemented with PTU (0.4% buffered tricaine or 1× tricaine). For injection to the circulation, 2 dpf zebrafish were anesthetized in 0.4% buffered tricaine and oriented in a lateral orientation on an agarose bed. A volume of 2-5 nL of the cell or cell-bead mixture suspension at 1 million cells/20 µl (∼200 cells) were injected directly in the circulation *via* the posterior cardinal vein using a pulled micropipette. Fish from a given clutch were randomly divided into experimental groups prior to injection. Injected fish were screened between 4-16 hr. post-injection to check for successful dissemination of cells through the circulatory system specifically at brain region. For drug-treated cancer cells (CellTracker Deep Red; Thermo Fisher Scientific, #C34565) and control/DMSO cells (CellTracker Blue CMAC; Thermo Fisher Scientific, #C2210), co-injection was performed after mixing into 1:1 ratio right before injection.

### Intravital Microscopy of Tracking Intravascular and Extravasated Cancer Cells

A Zeiss 780 LSM confocal microscope with a Zeiss 20× EC Plan-Apochromat, 0.8 NA objective was used to monitor the dynamics of cancer cells at zebrafish brain based on one-photon confocal 12-bit images with a digital zoom of 0.6, resulting in a field of view of 707.1 µm x 707.1 µm for each tile of the image at the resolution of 1.38 µm per pixel for each tile of the image. Fish were simultaneously excited with 488 nm light from an argon laser with a total power of 25 mW, 561 nm light from a solid-state laser with a total power of 20 mW, and 633 nm light from a HeNe633 solid state laser with a total power of 5 mW while also recording transmittance. Images were taken on two tracks to minimize signal overlap. All lasers were set at 2% total power. The master gain for all channels was at or below 650. A beam splitter, MBS 488/561/633, was employed in the emission pathway to delineate the red (band-pass filters 585–640 nm), green (band-pass filters 500-570 nm), and far red (bandpass filters 650-740 nm) channels. Pinhole diameter was set at 90 µm. The zebrafish larvae were maintained at 33 °C during whole imaging sessions.

Zebrafish larvae were anesthetized and immobilized in a lateral orientation in 1.25% (w/v) low gelling temperature agarose (Millipore Sigma, #A9414-25G) dissolved in fish water. To enable high-resolution confocal imaging of mounted larvae, fish were laterally oriented in coverglass-bottom chamber slides (Thermo Fisher Scientific, #155383). PTU water supplemented with 0.1% (0.25× Tricaine) buffered tricaine was then added to the imaging chamber to keep the larvae anesthetized healthier over the course of the experiment.

### Optical Tweezer (OT) Based Active Microrheology Setup and Measurement

We used the OT instrument as previously described to measure cancer cell mechanics change over the course of extravasation based on *in vivo* (zebrafish) and *in vitro* (microfluidics) models [29, 36, 37, 73]. Our instrument consists of a 1064 nm trapping beam (IPG Photonics, #YLR-20-1064-Y11) and a 975 nm detection beam (Lumics, #LU0975M00-1002F10D). The trapping beam is oscillated by a dual axis acousto-optic deflector (AOD) (IntraAction, #DTD274HD6). An iris after the AOD selects the doubly diffracted beam (i.e. 1st order in both transverse axes). The AOD receives control signals from radio frequency generating cards (Analog Devices, #AD9854/PCBZ) with onboard temperature-controlled crystal oscillators (Anodyne Components, ZKG10A1N-60.000M). The cards are controlled by digital outputs from a data acquisition card (National Instruments, #PCIe-5871R FPGA). AODs are mounted on 5-axis adjustable mounts (Newport, New Focus 9081). Both beams are shuttered electronically (Uniblitz, #VS1452Z0R3). Polarizing beam splitter cubes (Thorlabs, PBS23) linearly polarize the trapping beam. Before entering the AOD, the beam is attenuated manually by half-wave plates (Thorlabs, WPH05M-1064) or electronically via analog output from the data acquisition card. To detect the displacement of the trapping beam’s position, a beam sampler mirror (Thorlabs, #BSF10-C) and neutral density (ND) filter (Thorlabs, #NENIR210B) after the AOD direct a small amount of power (∼1 %) onto the ‘trap’ quadrant photodiode (QPD) (First Sensor, #QP154-QHVSD). The trapping beam is expanded by a lens pair (Thorlabs, 100 mm – #LA1509-C; 200 mm – #AC508-200-B) and directed into the microscope (Nikon, Eclipse Ti-U) backport with a broadband mirror (Thorlabs, #400 BB1-EO3IR). A lens pair (Thorlabs, 50 mm – #LA1131-C; 200 mm – #AC508-200-B) expands the detection beam, which is then coupled into and aligned with the trapping beam by a dichroic mirror (Chroma, #T1020LPXR). A third lens pair (Thorlabs, 50 mm – #LA113-C; 125 mm – #LA1384-CA) expands both beams so the trapping beam slightly overfills the back aperture of the objective (Nikon, #MRDO7602 CFIPLAN-APO VC60XA WI 1.2 NA). A dichroic filter cube (Chroma, #ZT1064rdc-2p) sends both beams into the objective. A high numerical aperture (NA), long working distance (WD) condenser (Nikon, #WI 0.9NA) collects the light from the objective. Behind the condenser, a dichroic mirror (Chroma, #ZT1064RDC-2P) directs the detection beam through a relay lens that is positioned to image the back focal plane of the condenser onto the ‘detection’ QPD. The trapping beam is removed from the path to the QPD with a bandpass filter (Chroma, #ET980/20X). Time-correlated ‘trap’ and ‘detection’ QPD signals are collected by analog inputs of the DAQ card. Control and data collection are conducted in custom programs (National Instruments, LabVIEW). A charge-coupled device (CCD) camera is mounted to the optical table. This enables position adjustments in X, Y and Z to place the camera in a plane conjugate to the trapping beam AOD, back-aperture of the condenser, and detection QPD. The Hz-nm constant relating the AOD’s RF control signal (in Hz) to the beam displacement (∼nm) is calibrated by attenuating and focusing the beam on a coverslip and imaging the backscattered beam on the CCD camera. Before each experiment, the alignment of the beams and the back focal plane interferometer is confirmed. Laser power is measured at the microscope backport with a power meter (Fieldmate, Coherent) and adjusted to 200 mW at the half-wave plate. A flow chamber is constructed from a microscope slide and cover slip with double-sided tape (Scotch) and loaded by capillary action with latex beads (Life Technologies, #F13083) suspended at low concentration in water. A bead is trapped, and the trap is oscillated while simultaneously viewing the bead position in real time from the ‘detection’ QPD signal. The beam-coupling dichroic and the QPD position are adjusted until oscillations in both transverse axes are centered on the QPD. When the system is aligned, a thermal power spectrum is recorded and fitted to a Lorentzian to calculate the viscosity of the water to confirm the system is calibrated. Before measurements, the camera pixel coordinates of the trap’s position are found by fitting a centroid to the intensity of an image collected of a trapped, stationary bead in water.

With the bead positioned in the trap as described above, the trap is oscillated while both the ‘trap’ and ‘detection’ QPD signals are recorded. The oscillation is multiplexed, i.e. a superposition of sine waves of differing phase and frequency, with the same amplitude at each frequency (25.4 nm per frequency or as specified). The frequencies are prime numbers to avoid interference of harmonics with fundamentals. Four phases are interlaced to minimize the total amplitude of the composite waveform. This waveform is pulsed for 2s, followed by 2s with the trap stationary to record the bead’s passive motion. This is repeated until 7 active-passive pulse sequences are recorded. The optical trap stiffness is determined in situ from the active and passive power spectra of each measured bead while the trap is stationary, using the active-passive calibration method as described below. Together with the inverse position detection sensitivity, the trap stiffness, bead mass m and hydrodynamic radius a, the trajectories yield the complex modulus as a function of frequency, G*(ω), of each bead’s surrounding microenvironment. The complex modulus is calculated as (Eq. 1)

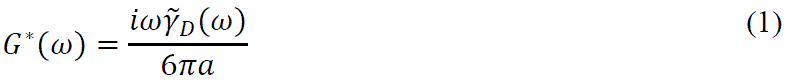

where a is the bead hydrodynamic radius and the friction relaxation spectrum, 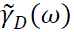, is shown as (Eq. 2)

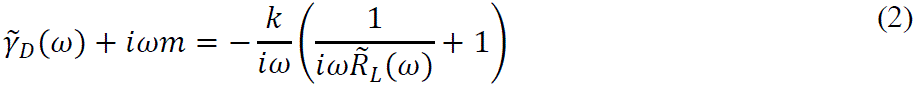

It relates to the active power spectrum, 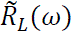, as (Eq. 3)

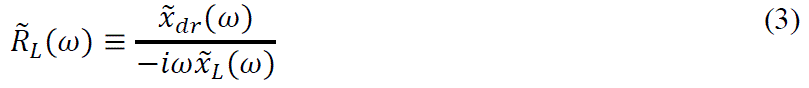

where 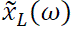 and 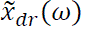 are as the Fourier transforms of the time series of the positions of the trapping laser and the driven bead respectively, recorded while the trap is oscillating

The stiffness (Eq. 4) is determined from the real part of the active power spectrum

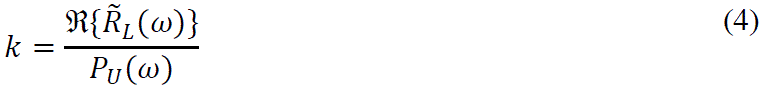

and the passive power spectrum (Eq. 5) where is the Fourier transform of the time series of the undriven bead’s thermally fluctuating position while the trap is held stationary.

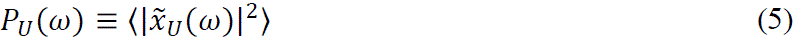

where 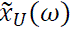 is the Fourier transform of the time series of the undriven bead’s thermally fluctuating position while the trap is held stationary.

Based on these equations, the complex modulus, G* = |G*|exp(iδ) = G′ + iG″, whose magnitude |G*| = (G′^2^ + G″^2^)^1/2^ shows the rigidity while G″/G′ encodes the hystersivity to determine liquid-like and solid-like characteristics. The Amplitude of oscillations was set to 20 nm per frequency to obtain a total amplitude of 200nm for the trap oscillation amplitude. For all amplitudes, laser power was set to 200 mW at the microscope back port. All OT experiments were controlled using custom LabVIEW programs. Data were analyzed by custom MATLAB programs.

## Microfluidic devices

Molds for microfluidic devices were fabricated using standard photolithography procedures[74]. AutoCAD (Autodesk, San Rafael, CA) was used to design five-inch by five-inch photolithography masks, which were purchased from Front Range Photomask (Lake Havasu City, AZ). To generate molds, four-inch silicon wafers (NOVA Electronic Materials, Flower Mound, TX) were dehydrated for 10 minutes on a 200 °C hotplate. SU-8 photoresist (Kayaku, Westborough, MA) layers were spun on wafers using a single wafer spin coater (Model WS-650S-8NPP/LITE, Laurell Technologies, North Wales, PA). An adhesion layer of SU-8 2000.5 was spun on the wafer, soft baked, flood-exposed with a mask aligner (Hybralign 200, Optical Associates, Inc., San Jose, CA), and post-exposure baked. The microchannel layer was then spun on the wafer to achieve an 8.39 μm thickness using SU-8 2007, as indicated by layer heights from the manufacturer. After soft baking, microchannel patterns were transferred to the wafers *via* UV exposure through the photomask, following a postexposure bake. The large 8.4 μm tall fluidic channels were patterned with SU-8 2050. Edge beads were removed prior to soft baking and the wafers were UV-exposed through the secondary photomask aligned to the first microchannel layer. During exposure of both layers, a long-pass filter (PL-360-LP, Omega Optical, Brattleboro, VT) was used. Patterned wafers were post-exposure baked, developed in SU 8 developer then rinsed with isopropanol and water. To enhance the mechanical properties of the SU 8 features, the wafers were hard-baked at 200 °C for approximately 5 minutes before ramping down to room temperature at 150°C/hr. The feature heights were measured using an optical profilometer (Filmetrics, San Diego, CA).

Final microfluidic devices were formed using replica molding with polydimethylsiloxane (PDMS; Sylgard 184 kit, Dow Corning, Midland, MI, USA) at a ratio of 10 parts prepolymer to 1-part crosslinker. PDMS devices were polymerized at 80 °C for ∼2 hours following degassing, and devices were then removed from the master mold. Devices were diced, and fluid access ports were punched with 4 mm biopsy punches (Miltex 69031-05, Electron Microscopy Sciences, Hatfield, PA).

For optical trap experiments, PDMS devices and two-well coverglass-bottom chamber slides (Nunc Lab-Tek Chambered #1.0 Borosilicate Coverglass slides: Cat. No. 155380, ThermoFisher Scientific, or ibidi μ-Slide 2 Well Glass Bottom #1.5H slides; Cat. No. 80287, ibidi) or glass bottom dishes (PELCO Black Wall Glass Bottom Dishes, glass 40 mm; Cat. No. 14035-120, Ted Pella Inc.) were exposed to oxygen plasma for 45 seconds using a PE-100 (Plasma Etch) at 150W. PDMS devices were then irreversibly bound to the slides and filled with PBS. Device walls were coated with 1% BSA (Millipore Sigma, Cat. No. A7906-100G) diluted in PBS by adsorption for 1 h at 37°C, 5% CO_2_. Devices were washed with PBS in triplicate following removal of the BSA solution.

Cells were detached with 10 mM EDTA in PBS containing 1 μg/ml Hoechst 33342 (Cat. No. H3570, ThermoFisher Scientific) and 1 μM CellTracker Deep Red (Cat. No. C34565, ThermoFisher Scientific) for 20 min at 37°C, 5% CO_2_. Cells were resuspended in growth medium, centrifuged for 5 min at 1000 rpm, resuspended in growth medium, counted, and resuspended in growth medium to a concentration of 1.5 million cells per ml. The cell suspension was incubated with 1 μm-diameter carboxylate-modified microspheres (FluoSpheres Carboxylate-Modified Microspheres, 1.0 μm, yellow-green fluorescent, Cat. No. F8823, ThermoFisher Scientific) at a ratio of 10 μL cell suspension per 1 μL microsphere stock (2% solids) for 30 minutes at 37°C under gentle rotation. The suspension was centrifuged for 3 min at 300 rcf, resuspended in 1 ml of medium, and centrifuged again for 3 min at 300 rcf. The pellet was resuspended in full growth medium to a concentration of 2×10^6^ cells/ml. Cells were introduced to microfluidic devices by flow via the addition of 20 μL (straight microchannels) or 10 μL (dual stage microchannels) of cell suspension. After seeding for approximately 30 minutes, the cell suspension was removed, and wells were filled with full growth medium. Cells migrated in the absence of flow or chemical gradients.

To obtain timelapse images of migrating cells, devices were mounted in a stage top incubator on a Zeiss 780 LSM confocal microscope. Devices were maintained at 37°C, 5% CO_2_ and imaged every 10 min for up to 17 h. A Zeiss 20× EC Plan-Apochromat, 0.8 NA objective and a digital zoom of 0.8 was used to capture 12-bit images with a field of view of 531.37 μm x 531.37 μm (512 pixels x 512 pixels) with pixel dwell times of 1.58 ms. Multiple positions were used to image the microchannels of each device. Samples were excited with 405 nm light from a diode laser with a total power of 30 mW and 633 nm light from a HeNe633 solid state laser with a total power of 5 mW. Lasers were used at 2% of total power. Two beam splitters, MBS 488/561/633 and MBS 405, were employed in the emission pathway to collect blue (band-pass filters 410-483 nm), green (band-pass filters 508-526 nm), and far red (band-pass filters 638-647 nm) light. Transmitted light was also collected. Pinhole diameter was set at 100 μm. Gain was set at or below 700 for each channel.

### Real-time Deformability Cytometry (RT-DC)

Cells of interest were detached as described above. Then, cells were immediately resuspended in 1000 µl RT-DC measurement buffer, (Zellmechanik Dresden, Cell Carrier), at room temperature and measured after 1 hr. based on a published protocol[44]. Right before RT-DC measurements, cells in CellCarrier were filtered *via* cell strainer 40µm to dissociate any cell clumps. Cells were passed through a microfluidic channel constriction of 30µm x 30µm in cross-section (Zellmechanik Dresden, Flic30) by applying a constant flow rate (0.16µl/s, 0.24µl/s, and 0.32µl/s) into a channel to induce deformation of cells. A high-speed complementary metal-oxide semiconductor (CMOS) camera equipped with *The AcCellerator* module (Zellmechanik Dresden) captures images of all flowing cells at the channel to yield cell deformability, cell area (µm^2^), and Young’s modulus (kPa)[41]. Each measurement was controlled by the acquisition software Shape-In 2 (Zellmechanik Dresden). Additionally, the recorded data was exported *via* analysis software Shape-Out 2 (Zellmechanik Dresden) for further data processing through Prism programs for curation of graphs and statistical analysis. Approximate fluid stress calculation is shown in Supplementary Table 1.

### Drug assay (YAP inhibition)

Verteporfin (Millipore Sigma, # SML0534-25MG) stock was prepared as 10 mg/ml in DMSO and diluted as needed for the drug treatment assays. 100k cells were seeded in a well of a 6 well plate for 24 hours to prepare 3 complete well plates (total 18 wells total). Then, a well from each plate was changed to fresh media with 0.1% v/v DMSO as a vehicle control while the remaining wells were filled with fresh media containing drug at final concentration between 0.1 and 10 µg/ml. The cells were placed in the incubator for another 24 hours and then detached with 10 mM EDTA in PBS (as described above). Cells were counted by cell counter to quantify total number of cells and number of dead cells.

### Immunofluorescence to quantitate YAP Nuclear-translocation

1 x 10^5^ cells from control (0.1% DMSO) and drug-treated cells were seeded after cell detachment into a 4 well plate (two each) (ibidi, #80426) with fresh growing media without drug or DMSO. After ∼16 hours, cells were fixed for 10 minutes at room temperature (RT) in 4% paraformaldehyde/1×PBS (diluted from a 16% paraformaldehyde). Fixed cells were washed five times with 1×PBS for 5 minutes each. Washed cells were then permeabilized with 0.1% Triton X-100/1×PBS for 10 minutes at RT. Permeabilized cells were washed five times with 1×PBS for 5 minutes each. Then, cells were blocked with 5% goat serum, 0.3% Triton X-100, 1×PBS for 1 hour. Blocked cells were washed five times with 1×PBS for 5 minutes each. Cells were then incubated in a 1:200 dilution of rabbit monoclonal anti-YAP antibody (Cell Signaling Technology, #14074S) with 1% goat serum, 0.3% Triton X-100, 1×PBS for 1 hr. at RT to visualize YAP. Stained cells were washed in 1×PBS five times for 5 minutes each. Antibody-stained cells were then transferred to a 1:400 secondary antibody cocktail containing an AlexaFluor Plus 488 goat anti-rabbit secondary antibody (Thermo Fisher Scientific, #A-11008) and a 1:40 AlexaFluor Phalloidin (A22284) in 1% goat serum, 0.3% Triton X-100, 1×PBS for 30-60 minutes at RT in dark. Cells were washed in 1×PBS for five minutes each before staining nucleus with DAPI (4’,6-Diamidino-2-Phenylindole, Dihydrochloride; Thermo Fisher Scientific, #D1306) in 1×PBS for 5 minutes. Nucleus-stained cells were washed in 1×PBS for 5 minutes before imaging with a Zeiss 780 LSM confocal microscopy.

### YAP-inhibition of human patient derived samples

Patient tumor samples were shipped overnight in DMEM on ice. The samples were dissociated using the Miltenyi Biotec gentleMACS Octo Dissociator with heater and the Miltenyi Biotec tumor dissociation kit, human using the manufacturer’s protocol. Briefly, once the fresh tumor specimens arrived, the samples were cut up using a scalpel. The sample was then placed into a “C tube” with 4.7mL DMEM, 200uL Enzyme H, 100uL Enzyme R, 25uL Enzyme A. The “C tube” was placed onto the gentleMACS Dissociator and program 37C_h_TDK_1 was run. The sample was strained using MACS SmartStrainer 70um. The strainer was washed with 20mL DMEM. The cells were centrifuged at 300×g for 7 minutes and resuspended into FBS and 10%DMSO and frozen at –80C.

Right before experiments, samples were then thawed and rinsed with a serum-free conditioned media. A serum-free conditioned medium was comprised of DMEM/F12 supplemented with 4 μg/mL BSA, 5ng/mL insulin, 100 U/mL penicillin, and 100 μg/mL streptomycin, 20 ng/mL recombinant epidermal growth factor (EGF), 20 ng/mL basic fibroblast growth factor (bFGF), and 1X B27TM supplement. Then, the patient samples were treated with control/DMSO and verteporfin (YAP inhibition) at 1μg/ml in suspension for 4 hours in a cell culture incubator before running for RTDC. For patient samples, RTDC was run at channel width of 20 μm in CellCarrierB (6.21*10^-3^ Pa*s).

### Immunoblot quantitation of YAP and Lamin A/C activity

Protein lysates from control/DMSO and VP (YAP inhibitor) were separated by electrophoresis on 4-12% bis-Tris SDS PAGE gels, with MOPS buffer and transferred to a nitrocellulose membrane. After blocking, the membranes were incubated overnight for 10 hours at 4°C with the primary antibodies: Rabbit Polyclonal anti-phospho YAP (ser127) (Cell Signaling, #4911S) at 1:1000 dilution or Rabbit Monoclonal anti-phospho Lamin A/C (Ser22) (D2B2E) (Cell Signaling, #13448S) at 1:1000, Rabbit Monoclonal anti-YAP (D8H1X) XP® (Cell Signaling, #14074S at 1:1000, Mouse Monoclonal anti-Lamin A/C (4C11) (Cell Signaling, #4777S) at 1:1000, and Rabbit Monoclonal anti-GAPDH (Cell Signaling, #2118S) at 1:1000. These primary antibodies were detected using Polyclonal Goat anti-Rabbit coupled to IRDye 680RD (LI-COR Bioscences, #925-68071) at 1:10000 or Polyclonal Goat anti-Mouse coupled to IRDye 680RD (LI-COR, #925-68070) at 1:10000.

### Mass Spectrometry

For proteomic analysis of YAP inhibition (verteporfin), control and drug treated MDA-MB-231 cells were washed with PBS and then detached as described above. Then, 2 million cells from each different condition were mixed with 1 ml of lysis buffer containing 1 part protease inhibitor cocktail set 1 (Calbiochem, #539131), 1 part phosphatase inhibitor cocktail set 2 (Millipore Sigma, #P5726), 1 part phosphatase inhibitor cocktail set 3 (Millipore Sigma, #P0044), and 100 parts M-PER Mammalian Protein Extraction Reagent (Thermo Fisher Scientific, #78501) to prepare protein lysates. The lysates were sonicated for 20 seconds and spun down 5 minutes at 1600 rpm. The supernatants were stored at –80 °C prior to protein digestion and mass spectrometry. Lysates were prepared on three different days.

For in-solution digestion of total proteins, frozen lysates were thawed, and protein concentrations measured using a BCA Protein Assay Kit (Thermo Scientific, #23227). Between 200-250 µg of lysates were reduced with 10 mM DTT at 56°C and alkylated with 20 mM iodoacetamide at 25°C in the dark. The samples were then digested with trypsin (50:1 protein: enzyme) overnight. The digested peptides were desalted using C18 spin desalting columns (Thermo Scientific, #89852) and lyophilized. The dried peptides were suspended in triethylammonium bicarbonate (TEAB), and concentrations were measured using colorimetric BCA peptide assay (Thermo Scientific, #23275). Next, 100 µg of digested peptides were labeled with the TMT six-plex reagent (Thermo Scientific, #90061) following the manufacturer’s protocol. Following labeling, peptide samples were pooled, lyophilized, and fractionated using high pH reversed-phase peptide fractionation (Thermo Scientific, #84868) following manufacturer’s instructions. Fractions were analyzed by mass spectrometry on an Orbitrap Exploris 480 (Thermo Electron) mass spectrometer. The peptides were separated on a 75 µm x 25 cm, 2 µm PepMap reverse phase column (Thermo) at 300 nL/min using an UltiMate 3000 RSLCnano HPLC (Thermo) and eluted directly into the mass spectrometer. For analysis, parent full-scan mass spectra were acquired at 120,000 FWHM resolution and product ion spectra at 45,000 resolution with a 0.7 m/z isolation window. Proteome Discoverer 3.0 (Thermo) was used to search the data against the human database from Uniprot using SequestHT. The search was limited to tryptic peptides, with maximally two missed cleavages allowed. Cysteine carbamidomethylation and TMT modification of lysine and peptide N-termini were set as fixed modifications, with methionine oxidation as a variable modification. The precursor mass tolerance was 10 ppm, and the fragment mass tolerance was 0.02 Da. The Percolator node was used to score and rank peptide matches using a 1% false discovery rate. TMT quantitation was performed using the Reporter Ions Quantifier nodes with correction of the values for lot-specific TMT reagent isotopic impurities.

Then, ingenuity Pathway Analysis software (IPA, Qiagen) was used for pathway analysis. Using the list of significantly (p < 0.05) differentially expressed proteins, the Canonical Pathway analysis, Disease & Function analysis, and networks analysis were performed by IPA.

### Data Analysis Processing and Statistics

Addition to analyzed OT data via custom MATLAB, data and graphs were plotted using Prism programs. Schemes (BioRender) and graphs were assembled for the figures using Adobe Illustrator. Statistical analysis was carried out in Prism. Details on statistical tests are given in each figure legend.

## Figures and Figure Legends

**Supplementary Figure 1:**
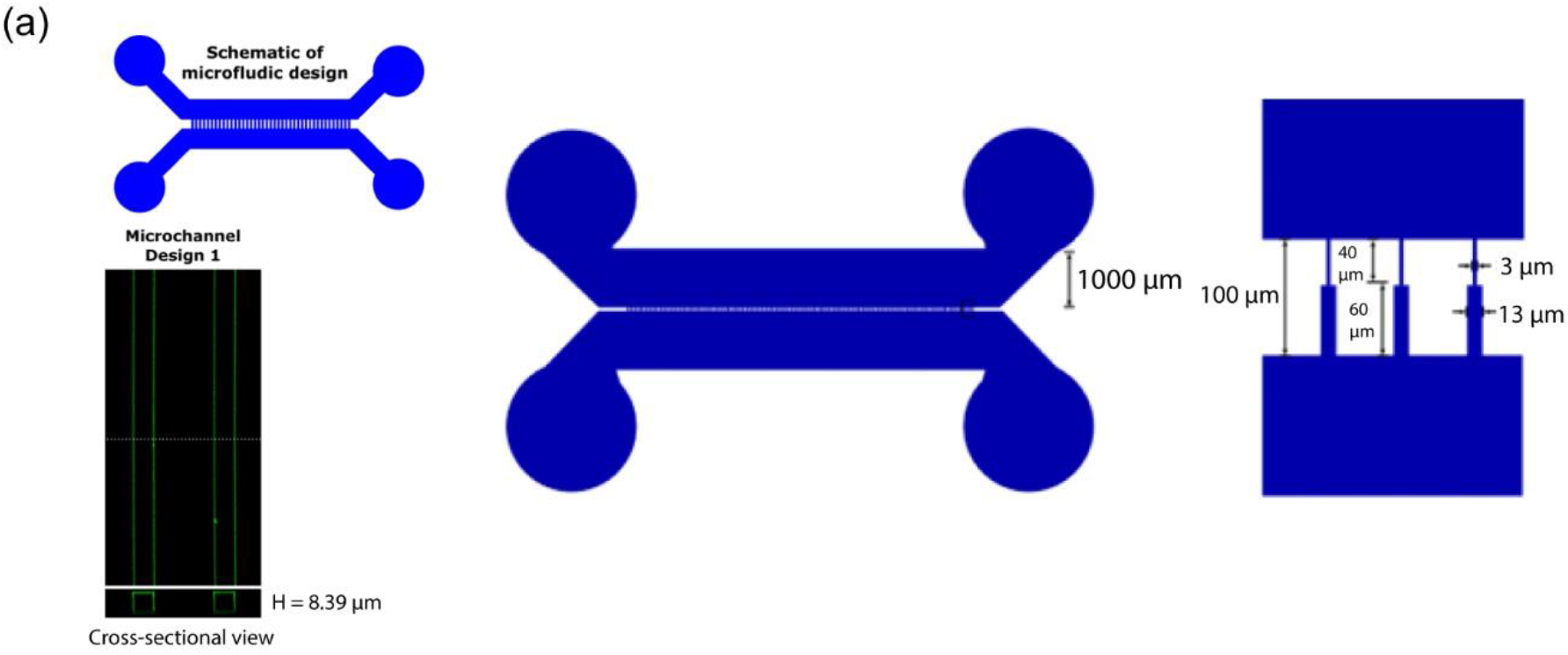
(a) Scheme of extravasation mimicking device design

**Supplementary Figure 2:**
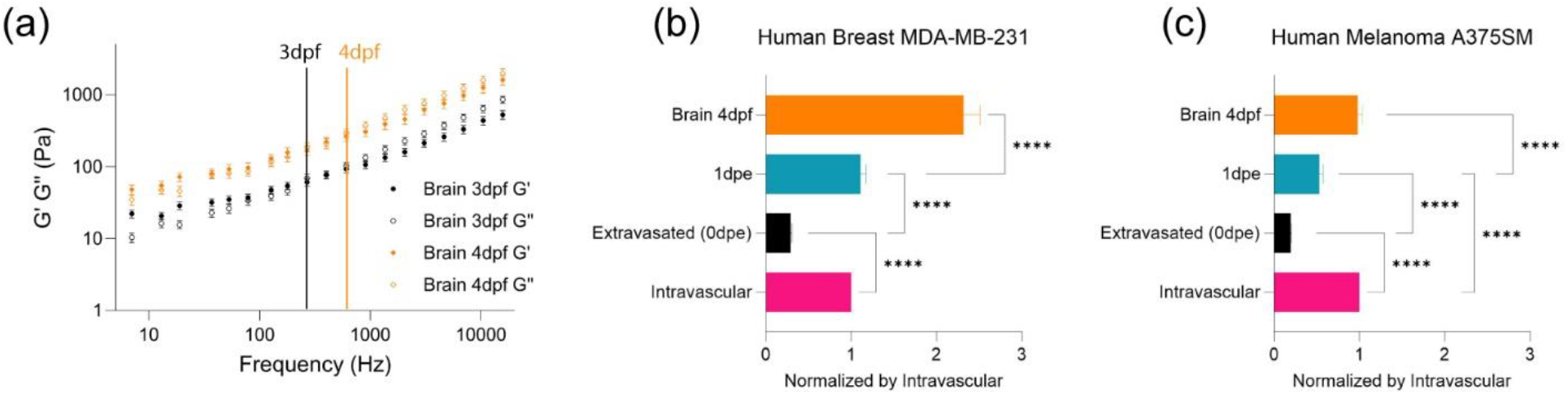
(a) log-log plot of *in vivo* brain tissue mechanics (elastic modulus, G’, and viscous modulus, G’’) and frequencies (7Hz to 15kHz) in the function of ‘3dpf’ (number of fish = 5, n = 132) and ‘4dpf’ (number of fish = 5, n = 154). Error bars in standard of error. Crossover frequency is assigned for each condition to point out the frequency where G’’ becomes more dominant than G’. (b) Normalized bar graph of complex modulus for ‘Breast (Intravascular)’, ‘Breast (0dpe)’, ‘Breast (1dpe)’, and ‘Brain 4dpf’ in respect to ‘Breast (Intravascular)’ based on 19 different frequencies from 7Hz to 15kHz. **** p<0.0001, paired two-tailed t-tests. (c) Normalized bar graph of complex modulus for ‘Melanoma (Intravascular)’, ‘Melanoma (0dpe)’, ‘Melanoma (1dpe)’, and ‘Brain 4dpf’ in respect to ‘Melanoma (Intravascular)’ based on 19 different frequencies from 7Hz to 15kHz. **** p<0.0001, paired two-tailed t-tests.

**Supplementary Figure 3:**
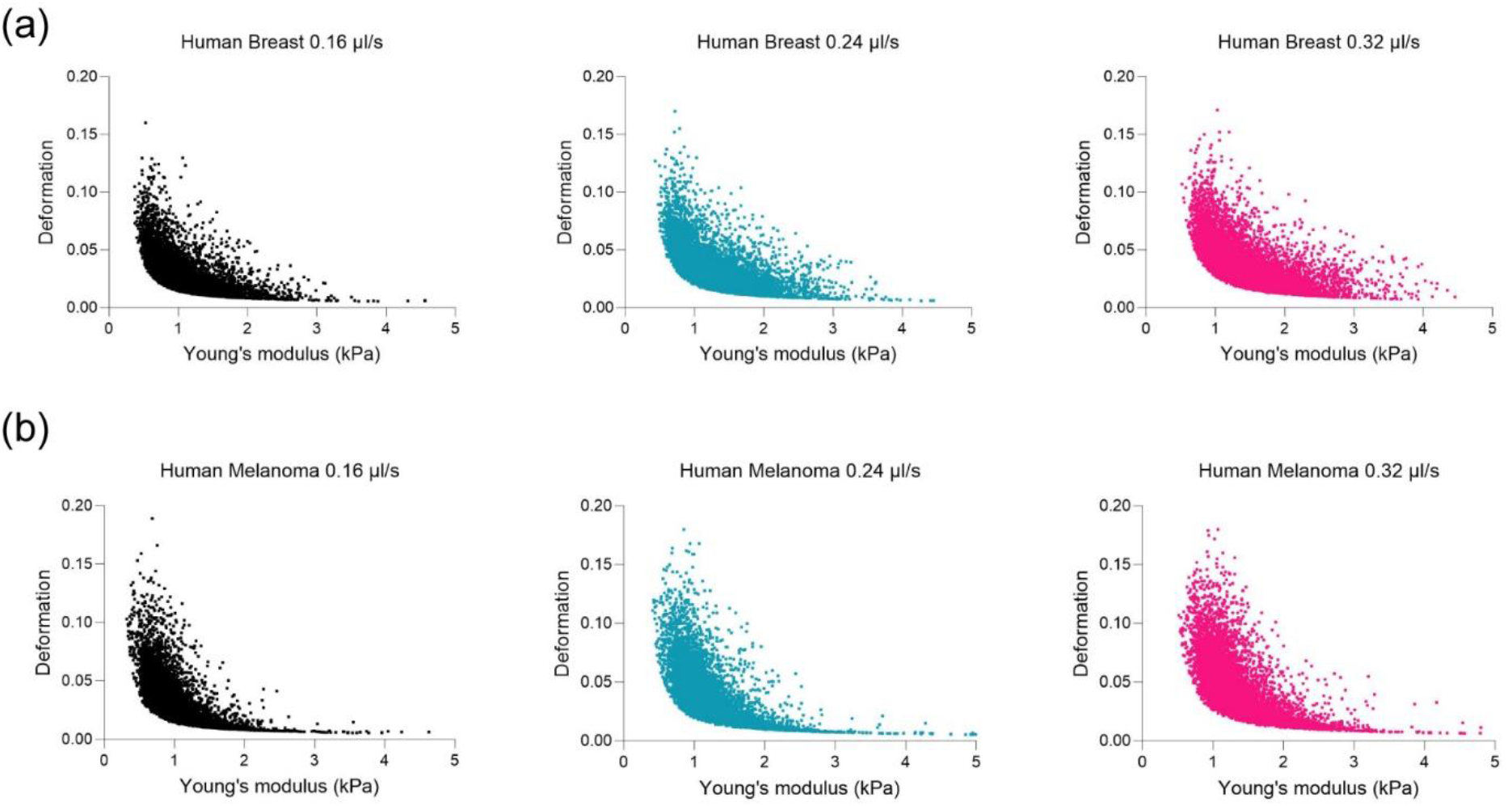
(a) Deformation and Young’s modulus (kPa) plot of human breast cancer cell at different flow velocities, 0.16 µl/s (n=4906), 0.24 µl/s (n=7398), and 0.32 µl/s (n=9506) (b) Deformation and Young’s modulus (kPa) of human melanoma at different flow velocities, 0.16 µl/s (n= 3954), 0.24 µl/s (n= 6284), and 0.32 µl/s (n= 8442).

**Supplementary Figure 4:**
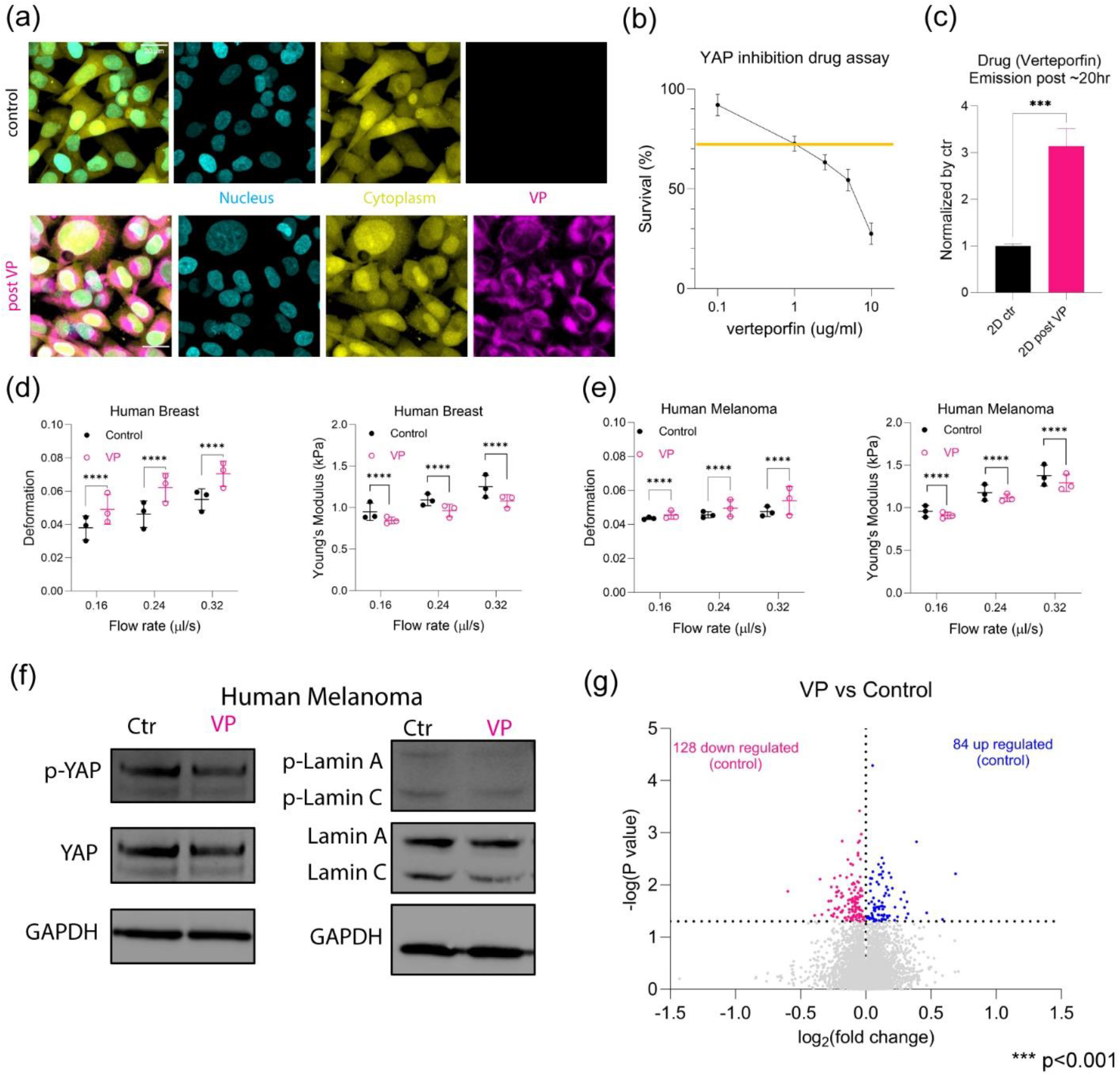
(a) Live imaging of cells between control/DMSO and verteporfin (VP) treated human breast cancer cells (b) YAP inhibition (VP) drug assay from 0.01 to 10 µg/ml (N=3) (c) Emission intensity of VP between control/DMSO cells and post VP treated cells (after ∼20hr) (N=3). *** p<0.001, paired two-tailed t-tests. (d) Deformation comparison for human breast cancer cells between control/DMSO (N=3; per each replicate more than 1000 cells) and VP (N=3; per each replicate more than 1000 cells) at 0.16 µl/s, 0.24 µl/s, and 0.32 µl/s. **** p<0.0001, paired two-tailed t-tests. Young’s modulus (kPa) comparison for human breast cancer cells between control/DMSO (N=3; per each replicate more than 1000 cells) and VP (N=3; per each replicate more than 1000 cells) at 0.16 µl/s, 0.24 µl/s, and 0.32 µl/s. **** p<0.0001, paired two-tailed t-tests. (e) Deformation comparison for human melanoma between control/DMSO (N=3; per each replicate more than 1000 cells) and VP (N=3; per each replicate more than 1000 cells) at 0.16 µl/s, 0.24 µl/s, and 0.32 µl/s. **** p<0.0001, paired two-tailed t-tests. Young’s modulus (kPa) comparison for human melanoma between control/DMSO (N=3; per each replicate more than 1000 cells) and VP (N=3; per each replicate more than 1000 cells) at 0.16 µl/s, 0.24 µl/s, and 0.32 µl/s. **** p<0.0001, paired two-tailed t-tests. (f) Representative western blot of total YAP with phosphorylated YAP and total Lamin A/C with phosphorylated Lamin A/C for human melanoma (N=3). (g) Volcano plots, showing up-regulated and down-regulated proteins for control/DMSO.

**Supplementary Figure 5:**
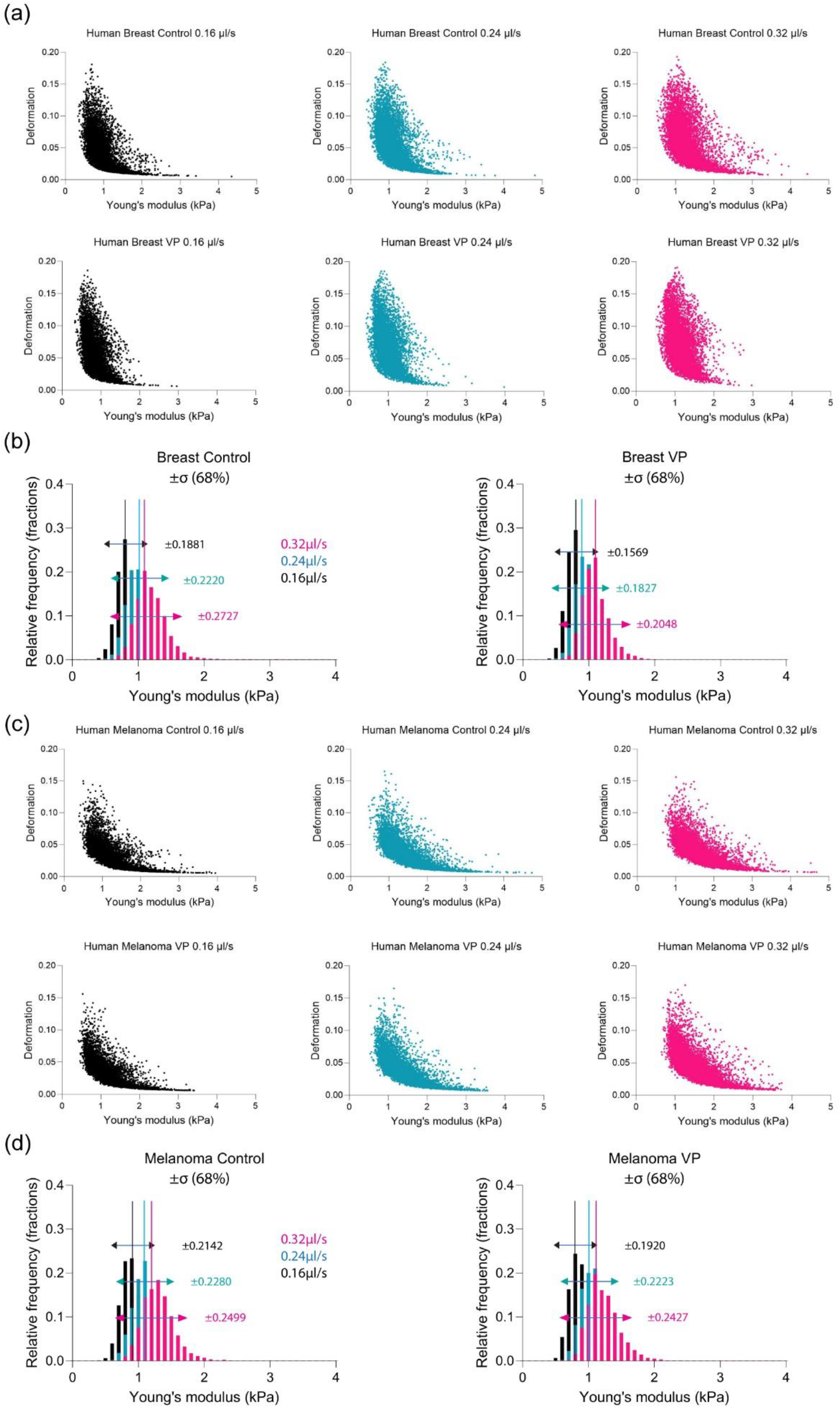
(a) Deformation and young’s modulus (kPa) plot of human breast cancer cell control/DMSO at different flow velocities, 0.16 µl/s (n=6495), 0.24 µl/s (n=8401), and 0.32 µl/s (n=8712) with a fit based on ‘deformation=(slope)/young’s modulus. Deformation and young’s modulus (kPa) plot of human breast cancer cell VP at different flow velocities, 0.16 µl/s (n=8734), 0.24 µl/s (n=10095), and 0.32 µl/s (n=10637) with a fit based on ‘deformation=(slope)/young’s modulus. (b) Normalized histogram of human breast cancer cell for young’s modulus (kPa) between control/DMSO and VP at different flow velocities, 0.16 µl/s (control n=6495, VP n=8734), 0.24 µl/s (control n=8401, VP n=10095), and 0.32 µl/s (control n=8712, VP n=10637), with standard deviation (σ, ±68%). (c) Deformation and young’s modulus (kPa) plot of human melanoma control at different flow velocities, 0.16 µl/s (n=9979), 0.24 µl/s (n=9991), and 0.32 µl/s (n=9928) with a fit based on ‘deformation=(slope)/young’s modulus. Deformation and young’s modulus (kPa) plot of human melanoma VP at different flow velocities, 0.16 µl/s (n=9992), 0.24 µl/s (n=9991), and 0.32 µl/s (n=9988) with a fit based on ‘deformation=(slope)/young’s modulus. (d) Normalized histogram of human melanoma for young’s modulus (kPa) between control/DMSO and VP at different flow velocities, 0.16 µl/s (control n=9979, VP n=9992), 0.24 µl/s (control n=9991, VP n=9991), and 0.32 µl/s (control n=9928, VP n=9988), with standard deviation (σ, ±68%).

**Supplementary Table 1:**
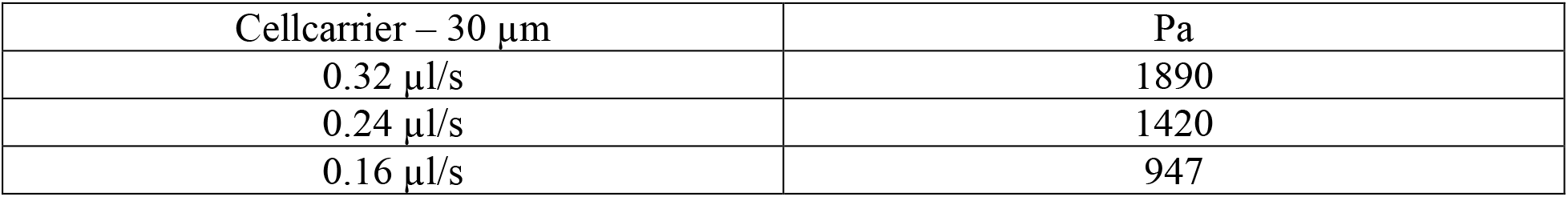
Calculated stress for channel width of 30 µm and Cellcarrier solution.

| Cellcarrier – 30 $\mu\text{m}$ | Pa |
| --- | --- |
| 0.32 $\mu\text{l/s}$ | 1890 |
| 0.24 $\mu\text{l/s}$ | 1420 |
| 0.16 $\mu\text{l/s}$ | 947 |

Stress under laminar flow based on Poiseuille’s law

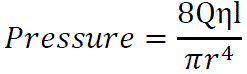

Where Q is flow velocity, η is viscosity of fluid, l is the channel length, r is radius of channel CellCarrier = 3.92*10^-3^ (Pa*s)

**Supplementary Movie 1**: Extravasation moment of human breast cancer cells (MDA-MB-231) at zebrafish brain after injection of cells into the circulation of zebrafish, Tg(flk:mCherry/MRC1a:EGFP), at 2dpf. Blood vessel (flk) is in magenta, vein (MRC1a) is in yellow, and human breast cancer cells are in blue.

**Supplementary Movie 2**: Human breast cancer cells (MDA-MB-231) with bead migrating in narrow channel inside of microfluidic device.

**Supplementary Movie 3**: Tracking extravasated human breast cancer cells (MDA-MB-231) at zebrafish brain right after extravasation from 3dpf/1dpi overnight to 4dpf/2dpi. Blood vessel (flk) and vein (MRC1a) are in grey while human breast cancer cells are in blue.

**Supplementary Movie 4**: Live cell imaging of control/DMSO cancer cells with nucleus (H2B) in magenta and membrane dye (CellTracker) in green inside of extravasation mimicking microfluidics devices at different time interval.

**Supplementary Movie 5**: Live cell imaging of VP (Yap inhibition) treated cancer cells with nucleus (H2B) in magenta and membrane dye (CellTracker) in green inside of extravasation mimicking microfluidics devices at different time interval.

**Supplementary Movie 6**: Overnight movie after co-injecting control/DMSO (cyan) and VP (magenta) treated human breast cancer cells into circulation at 2dpf using Tg(flk:mCherry/MRC1a:EGFP); flk and MRC1a are labelled as grey.

## References

1. Wakefield, L., S. Agarwal, and K. Tanner, Preclinical models for drug discovery for metastatic disease. Cell, 2023. 186(8): p. 1792–1813.

2. Fidler, I.J., D.M. Gersten, and I.R. Hart, The biology of cancer invasion and metastasis. Adv Cancer Res, 1978. 28: p. 149–250.

3. Heidrich, I., B. Deitert, S. Werner, and K. Pantel, Liquid biopsy for monitoring of tumor dormancy and early detection of disease recurrence in solid tumors. Cancer Metastasis Rev, 2023. 42(1): p. 161–182.

4. Butler, T.P. and P.M. Gullino, Quantitation of Cell Shedding into Efferent Blood of Mammary Adenocarcinoma1. Cancer Research, 1975. 35(3): p. 512–516.

5. Ganesh, K. and J. Massagué, Targeting metastatic cancer. Nature Medicine, 2021. 27(1): p. 34–44.

6. Kalukula, Y., A.D. Stephens, J. Lammerding, and S. Gabriele, Mechanics and functional consequences of nuclear deformations. Nature Reviews Molecular Cell Biology, 2022. 23(9): p. 583–602.

7. Jain, N., J. Moeller, and V. Vogel, Mechanobiology of Macrophages: How Physical Factors Coregulate Macrophage Plasticity and Phagocytosis. Annu Rev Biomed Eng, 2019. 21: p. 267–297.

8. Zhang, X., et al., Unraveling the mechanobiology of immune cells. Curr Opin Biotechnol, 2020. 66: p. 236–245.

9. Wyckoff, J.B., et al., Direct visualization of macrophage-assisted tumor cell intravasation in mammary tumors. Cancer Res, 2007. 67(6): p. 2649–56.

10. Pegoraro, A.F., P. Janmey, and D.A. Weitz, Mechanical Properties of the Cytoskeleton and Cells. Cold Spring Harb Perspect Biol, 2017. 9(11).

11. Hayward, M.K., J.M. Muncie, and V.M. Weaver, Tissue mechanics in stem cell fate, development, and cancer. Dev Cell, 2021. 56(13): p. 1833–1847.

12. Bailey, M.H., et al., Comprehensive Characterization of Cancer Driver Genes and Mutations. Cell, 2018. 173(2): p. 371–385 e18.

13. Cheng, D.T., et al., Memorial Sloan Kettering-Integrated Mutation Profiling of Actionable Cancer Targets (MSK-IMPACT): A Hybridization Capture-Based Next-Generation Sequencing Clinical Assay for Solid Tumor Molecular Oncology. J Mol Diagn, 2015. 17(3): p. 251–64.

14. Han, L., et al., The Genomic Landscape and Clinical Relevance of A-to-I RNA Editing in Human Cancers. Cancer Cell, 2015. 28(4): p. 515–528.

15. Hsia, C.-R., D.P. Melters, and Y. Dalal, The Force is Strong with This Epigenome: Chromatin Structure and Mechanobiology. Journal of Molecular Biology, 2023. 435(11): p. 168019.

16. Yu, W., et al., Cancer cell mechanobiology: a new frontier for cancer research. Journal of the National Cancer Center, 2022. 2(1): p. 10–17.

17. Chaudhuri, P.K., B.C. Low, and C.T. Lim, Mechanobiology of Tumor Growth. Chemical Reviews, 2018. 118(14): p. 6499–6515.

18. Gabarra-Niecko, V., M.D. Schaller, and J.M. Dunty, FAK regulates biological processes important for the pathogenesis of cancer. Cancer Metastasis Rev, 2003. 22(4): p. 359–74.

19. Winkler, J., A. Abisoye-Ogunniyan, K.J. Metcalf, and Z. Werb, Concepts of extracellular matrix remodelling in tumour progression and metastasis. Nature Communications, 2020. 11(1): p. 5120.

20. Bissell, M.J., H.G. Hall, and G. Parry, How does the extracellular matrix direct gene expression? J Theor Biol, 1982. 99(1): p. 31–68.

21. Nelson, C.M. and M.J. Bissell, Of extracellular matrix, scaffolds, and signaling: tissue architecture regulates development, homeostasis, and cancer. Annu Rev Cell Dev Biol, 2006. 22: p. 287–309.

22. Massague, J., TGFbeta in Cancer. Cell, 2008. 134(2): p. 215–30.

23. Azubuike, U.F. and K. Tanner, Biophysical determinants of cancer organotropism. Trends Cancer, 2023. 9(3): p. 188–197.

24. Paul, C.D., et al., Tissue Architectural Cues Drive Organ Targeting of Tumor Cells in Zebrafish. Cell Syst, 2019. 9(2): p. 187–206 e16.

25. Follain, G., et al., Hemodynamic Forces Tune the Arrest, Adhesion, and Extravasation of Circulating Tumor Cells. Dev Cell, 2018. 45(1): p. 33–52 e12.

26. Xiao, J., et al., Identifying drivers of breast cancer metastasis in progressively invasive subpopulations of zebrafish-xenografted MDA-MB-231. Molecular Biomedicine, 2022. 3(1): p. 16.

27. Campbell, N.R., et al., Cooperation between melanoma cell states promotes metastasis through heterotypic cluster formation. Dev Cell, 2021. 56(20): p. 2808–2825 e10.

28. Stoletov, K., et al., Visualizing extravasation dynamics of metastatic tumor cells. Journal of Cell Science, 2010. 123(13): p. 2332–2341.

29. Blehm, B.H., A. Devine, J.R. Staunton, and K. Tanner, In vivo tissue has non-linear rheological behavior distinct from 3D biomimetic hydrogels, as determined by AMOTIV microscopy. Biomaterials, 2016. 83: p. 66–78.

30. Harlepp, S., F. Thalmann, G. Follain, and J.G. Goetz, Hemodynamic forces can be accurately measured in vivo with optical tweezers. Molecular Biology of the Cell, 2017. 28(23): p. 3252–3260.

31. Kienast, Y., et al., Real-time imaging reveals the single steps of brain metastasis formation. Nat Med, 2010. 16(1): p. 116–22.

32. Chen, M.B., J.A. Whisler, J.S. Jeon, and R.D. Kamm, Mechanisms of tumor cell extravasation in an in vitro microvascular network platform. Integrative Biology, 2013. 5(10): p. 1262–1271.

33. Strilic, B. and S. Offermanns, Intravascular survival and extravasation of tumor cells. Cancer cell, 2017. 32(3): p. 282–293.

34. Fischer, M. and K. Berg-Sørensen, Calibration of trapping force and response function of optical tweezers in viscoelastic media. Journal of Optics A: Pure and Applied Optics, 2007. 9(8): p. S239.

35. Fischer, M., et al., Active-passive calibration of optical tweezers in viscoelastic media. Review of Scientific Instruments, 2010. 81(1).

36. Staunton, J.R., B. Blehm, A. Devine, and K. Tanner, In situ calibration of position detection in an optical trap for active microrheology in viscous materials. Opt Express, 2017. 25(3): p. 1746–1761.

37. Staunton, J.R., W.Y. So, C.D. Paul, and K. Tanner, High-frequency microrheology in 3D reveals mismatch between cytoskeletal and extracellular matrix mechanics. Proc Natl Acad Sci U S A, 2019. 116(29): p. 14448–14455.

38. Lu, P., V.M. Weaver, and Z. Werb, The extracellular matrix: A dynamic niche in cancer progression. Journal of Cell Biology, 2012. 196(4): p. 395–406.

39. Sturgess, V., U.F. Azubuike, and K. Tanner, Vascular regulation of disseminated tumor cells during metastatic spread. Biophysics Reviews, 2023. 4(1).

40. Follain, G., et al., Fluids and their mechanics in tumour transit: shaping metastasis. Nature Reviews Cancer, 2020. 20(2): p. 107–124.

41. Otto, O., et al., Real-time deformability cytometry: on-the-fly cell mechanical phenotyping. Nature Methods, 2015. 12(3): p. 199–202.

42. Mietke, A., et al., Extracting Cell Stiffness from Real-Time Deformability Cytometry: Theory and Experiment. Biophysical Journal, 2015. 109(10): p. 2023–2036.

43. Mokbel, M., et al., Numerical Simulation of Real-Time Deformability Cytometry To Extract Cell Mechanical Properties. ACS Biomaterials Science & Engineering, 2017. 3(11): p. 2962–2973.

44. Toepfner, N., et al., Detection of human disease conditions by single-cell morpho-rheological phenotyping of blood. eLife, 2018. 7: p. e29213.

45. Dupont, S., et al., Role of YAP/TAZ in mechanotransduction. Nature, 2011. 474(7350): p. 179–183.

46. Lee, H.J., et al., Fluid shear stress activates YAP1 to promote cancer cell motility. Nature Communications, 2017. 8(1): p. 14122.

47. Vigneswaran, K., et al., YAP/TAZ Transcriptional Coactivators Create Therapeutic Vulnerability to Verteporfin in EGFR-mutant Glioblastoma. Clinical Cancer Research, 2021. 27(5): p. 1553–1569.

48. Jang, M., et al., Matrix stiffness epigenetically regulates the oncogenic activation of the Yes-associated protein in gastric cancer. Nature Biomedical Engineering, 2021. 5(1): p. 114–123.

49. Xu, W., K.M. Alpha, N.M. Zehrbach, and C.E. Turner, Paxillin promotes breast tumor collective cell invasion through maintenance of adherens junction integrity. Molecular Biology of the Cell, 2021. 33(2): p. ar14.

50. Xue, Q., et al., Lack of Paxillin phosphorylation promotes single-cell migration in vivo. Journal of Cell Biology, 2023. 222(3): p. e202206078.

51. Snigdha, K., et al., Hippo Signaling in Cancer: Lessons From Drosophila Models. Front Cell Dev Biol, 2019. 7: p. 85.

52. Yu, F.-X., B. Zhao, and K.-L. Guan, Hippo Pathway in Organ Size Control, Tissue Homeostasis, and Cancer. Cell, 2015. 163(4): p. 811–828.

53. Zanconato, F., M. Cordenonsi, and S. Piccolo, YAP/TAZ at the Roots of Cancer. Cancer Cell, 2016. 29(6): p. 783–803.

54. Lamar, J.M., et al., The Hippo pathway target, YAP, promotes metastasis through its TEAD-interaction domain. Proc Natl Acad Sci U S A, 2012. 109(37): p. E2441–50.

55. Rianna, C., M. Radmacher, and S. Kumar, Direct evidence that tumor cells soften when navigating confined spaces. Molecular Biology of the Cell, 2020. 31(16): p. 1726–1734.

56. Dahl, K.N., A.J. Ribeiro, and J. Lammerding, Nuclear shape, mechanics, and mechanotransduction. Circ Res, 2008. 102(11): p. 1307–18.

57. Nikolic, M., G. Scarcelli, and K. Tanner, Multimodal microscale mechanical mapping of cancer cells in complex microenvironments. Biophys J, 2022. 121(19): p. 3586–3599.

58. Roberts, A.B., et al., Tumor cell nuclei soften during transendothelial migration. Journal of Biomechanics, 2021. 121: p. 110400.

59. Au, S.H., et al., Clusters of Circulating Tumor Cells: a Biophysical and Technological Perspective. Curr Opin Biomed Eng, 2017. 3: p. 13–19.

60. Nia, H.T., L.L. Munn, and R.K. Jain, Physical traits of cancer. Science, 2020. 370(6516).

61. Malek, A.M., S.L. Alper, and S. Izumo, Hemodynamic shear stress and its role in atherosclerosis. JAMA, 1999. 282(21): p. 2035–42.

62. Lammerding, J., et al., Lamins A and C but not lamin B1 regulate nuclear mechanics. J Biol Chem, 2006. 281(35): p. 25768–80.

63. Swift, J., et al., Nuclear lamin-A scales with tissue stiffness and enhances matrix-directed differentiation. Science, 2013. 341(6149): p. 1240104.

64. Owens, D.J., et al. Lamin Mutations Cause Increased YAP Nuclear Entry in Muscle Stem Cells. Cells, 2020. 9, DOI: 10.3390/cells9040816.

65. Benjamin, D.C., et al., YAP Enhances Tumor Cell Dissemination by Promoting Intravascular Motility and Reentry into Systemic Circulation. Cancer Res, 2020. 80(18): p. 3867–3879.

66. Petrie, R.J., H.M. Harlin, L.I. Korsak, and K.M. Yamada, Activating the nuclear piston mechanism of 3D migration in tumor cells. J Cell Biol, 2017. 216(1): p. 93–100.

67. Petrie, R.J., H. Koo, and K.M. Yamada, Generation of compartmentalized pressure by a nuclear piston governs cell motility in a 3D matrix. Science, 2014. 345(6200): p. 1062–5.

68. Enzo, E., et al., Aerobic glycolysis tunes YAP/TAZ transcriptional activity. The EMBO Journal, 2015. 34(10): p. 1349–1370.

69. Koo, J.H. and K.L. Guan, Interplay between YAP/TAZ and Metabolism. Cell Metab, 2018. 28(2): p. 196–206.

70. Blehm, B.H., N. Jiang, Y. Kotobuki, and K. Tanner, Deconstructing the role of the ECM microenvironment on drug efficacy targeting MAPK signaling in a pre-clinical platform for cutaneous melanoma. Biomaterials, 2015. 56: p. 129–39.

71. Afasizheva, A., et al., Mitogen-activated protein kinase signaling causes malignant melanoma cells to differentially alter extracellular matrix biosynthesis to promote cell survival. BMC Cancer, 2016. 16: p. 186.

72. Hanson, R.L. and E. Batchelor, Coordination of MAPK and p53 dynamics in the cellular responses to DNA damage and oxidative stress. Molecular Systems Biology, 2022. 18(12): p. e11401.

73. Nikolic, M., G. Scarcelli, and K. Tanner, Multimodal microscale mechanical mapping of cancer cells in complex microenvironments. Biophys J, 2022.

74. Duffy, D.C., J.C. McDonald, O.J.A. Schueller, and G.M. Whitesides, Rapid Prototyping of Microfluidic Systems in Poly(dimethylsiloxane). Analytical Chemistry, 1998. 70(23): p. 4974–4984.

